# Recovery of highly contiguous genomes from complex terrestrial habitats reveals over 15,000 novel prokaryotic species and expands characterization of soil and sediment microbial communities

**DOI:** 10.1101/2024.12.19.629313

**Authors:** Mantas Sereika, Aaron James Mussig, Chenjing Jiang, Kalinka Sand Knudsen, Thomas Bygh Nymann Jensen, Francesca Petriglieri, Yu Yang, Vibeke Rudkjøbing Jørgensen, Francesco Delogu, Emil Aarre Sørensen, Per Halkjær Nielsen, Caitlin Margaret Singleton, Philip Hugenholtz, Mads Albertsen

## Abstract

Genomes are fundamental to understanding microbial ecology and evolution. The emergence of high-throughput, long-read DNA sequencing has enabled recovery of microbial genomes from environmental samples at scale. However, expanding the microbial genome catalogue of soils and sediments has been challenging due to the enormous complexity of these environments. Here, we performed deep, long-read Nanopore sequencing of 154 soil and sediment samples collected across Denmark and through an optimised bioinformatics pipeline, we recovered genomes of 15,314 novel microbial species, including 4,757 high-quality genomes. The recovered microbial genomes span 1,086 novel genera and provide the first high-quality reference genomes for 612 previously known genera, expanding the phylogenetic diversity of the prokaryotic tree of life by 8 %. The long-read assemblies also enabled the recovery of thousands of complete rRNA operons, biosynthetic gene clusters and CRISPR-Cas systems, all of which were underrepresented and highly fragmented in previous terrestrial genome catalogues. Furthermore, the incorporation of the recovered MAGs into public genome databases significantly improved species-level classification rates for soil and sediment metagenomic datasets, thereby enhancing terrestrial microbiome characterization. With this study, we demonstrate that long-read sequencing and optimised bioinformatics, allows cost-effective recovery of high-quality microbial genomes from highly complex ecosystems, which remain the largest untapped source of biodiversity for expanding genome databases and filling in the gaps of the tree of life.

## Introduction

The vast majority of microorganisms are predicted to be undiscovered^1^. Traditionally, achieving genomes of new microbial species involves isolating and cultivating the microorganisms, followed by sequencing^2^. While this method has successfully yielded thousands of novel genomes^3^, culturing can be labour-intensive and time-consuming^4^, and most microbes are estimated to be unsuitable for isolation^5^. In the past decade genome-centric metagenomics has emerged as an alternative and expedient means of characterising microbial diversity through recovery of metagenome-assembled genomes (MAGs)^6,7^. Despite potential issues of contamination and incompleteness (e.g. different microbial strains^8^ or species^9^), metagenomics allows large-scale recovery of novel genomes from uncultured microorganisms^10–12^. To date, the Genome Taxonomy Database (GTDB, release 220) comprises 113,104 prokaryotic species, of which 72.5% are represented exclusively by MAGs^13^, highlighting the current limitations in culture-based genomics. Therefore, MAGs will be indispensable to obtain genomic coverage of the estimated 2 to 4 million prokaryotic species inhabiting the biosphere^14^.

Soil has the potential to greatly increase the number of microbial species in the databases given its enormous microbial diversity^15^. However, this complexity also makes soil exceptionally challenging for MAG recovery^15^. Several attempts have been made to improve MAG recovery from soil, such as reducing the complexity of the sample through species enrichment^16^, cell sorting^17^ or deep short-read sequencing (e.g. over 100 Gbp to several Tbp of sequencing data)^18-21^. However, none of these approaches have resulted in cost-effective recovery of high-quality microbial genomes. Hence, developing a solution for efficient recovery of high-quality MAGs from soil and other microbially complex habitats is regarded as the “grand challenge” of metagenomics^22^.

In recent years, long-read sequencing has supercharged our ability to recover high-quality microbial genomes from medium complexity samples^23–25^. This has been complemented by development of bioinformatic methods that improve MAG recovery from challenging samples through the use of deep learning algorithms^21,26^ or additional binning features^27–31^. Therefore, multiple sequencing and bioinformatic approaches have now become available for tackling the “grand challenge” of soil metagenomics.

Here, we performed deep long-read Nanopore sequencing (∼approx. 100 Gbp/sample) of 154 complex environmental samples collected as part of the Microflora Danica project, which aims to genomically catalogue microbial diversity in Denmark^32^. By developing a bioinformatics workflow that uses state-of-the-art metagenomic binning tools, combined with multi-coverage and iterative binning, we obtained over 15,000 species-level MAGs. The great majority (97.9 %) of these MAGs represent either novel microbial genera or species and significantly expand the microbial tree of life.

## Results

### 1. High-throughput MAG recovery from soils and sediments

#### 1.1. Thousands of MAGs (hundreds per sample) recovered from terrestrial habitats

Of the 10,686 environmental samples collected during the Microflora Danica sampling campaign^32^, 154 samples (125 soil, 28 sediment, 1 water) from 15 distinct habitats (**Table S1, Figure S1**) were selected (see Methods for selection criteria) for deep long-read Nanopore sequencing (**Figure 1A**) in order to explore assembly performance across a wide breadth of sample types. A total of 14.4 Tbp long-read data was generated with a median of 94.9 Gbp and an interquartile range (IQR) of 56.3-133.1 Gbp (**Figure 1B**). The sequence reads had a median length of 6.1 kbp (IQR: 4.6-7.3) (**Figure 1C**) and assembled into a total of 295.7 Gbp of metagenomic contigs, with a median contig N50 of 79.8 kbp (IQR 45.8-110.1) per sample. The majority of reads were assembled into contigs, as a median 62.2 % (IQR: 53.1-69.8) of the sequence data was mapped back to the assemblies (**Figure 1D**).

**Figure 1:**
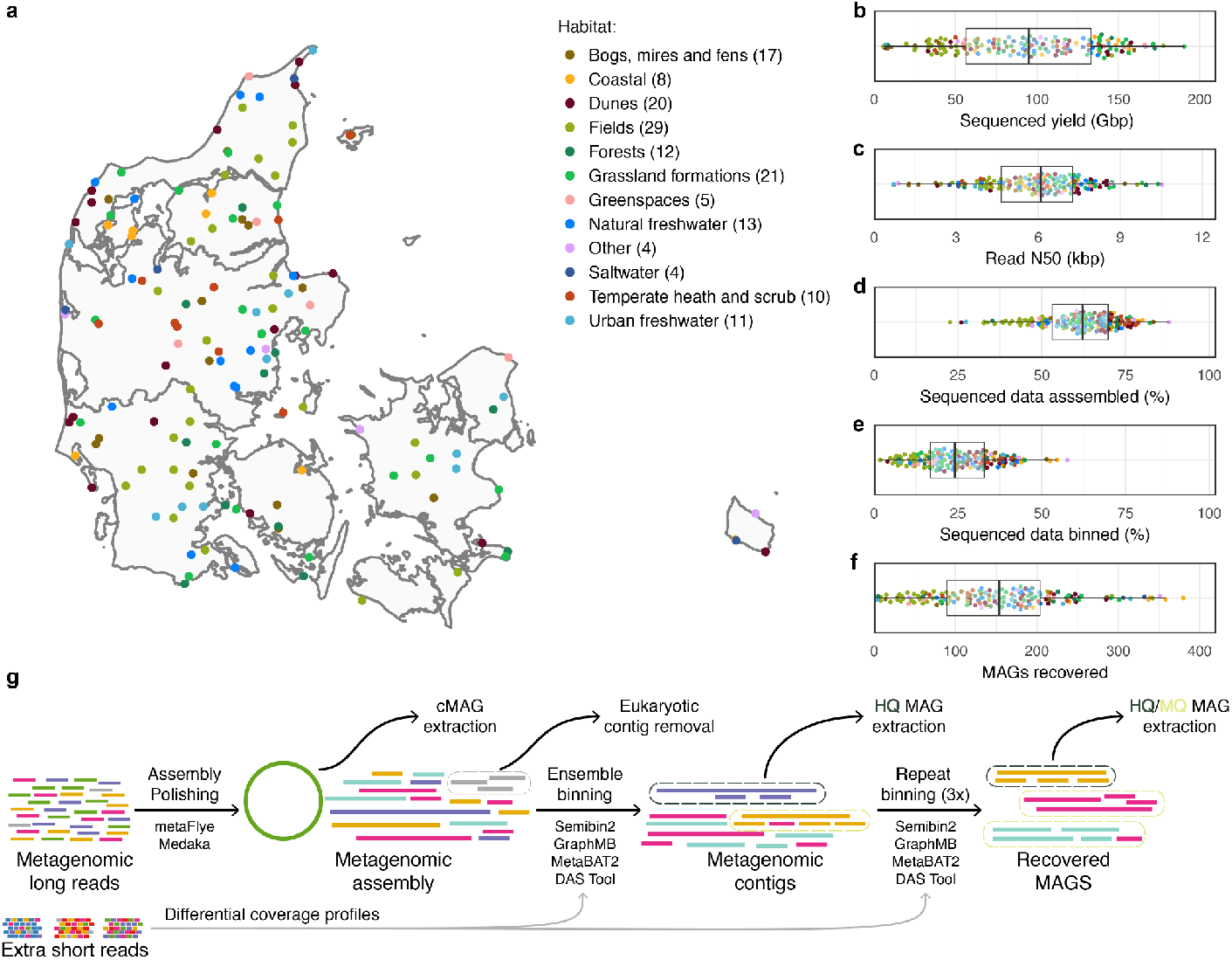
Overview of the sequenced environmental samples. **A)** Geographic distribution of the samples, coloured by sample habitat (Microflora Danica Ontology level 1). Samples from habitats, which occur once in the dataset, are grouped into the “Other” habitat category. **B)** Per-sample sequencing yield in Gbp. **C)** Per-sample sequenced read N50 values in kbp. **D)** Per-sample percentage of sequencing data that was estimated as assembled into contigs. **E)** Estimates for per-sample percentage of sequenced data represented by the recovered high or medium quality MAGs. **F)** Per-sample count for the recovered high or medium quality MAGs. **G)** Schematic overview of mmlong2 metagenomics workflow. “Extra short reads” refers to the shallow metagenome datasets from the Microflora Danica project^32^ used for differential coverage binning (see Methods). Abbreviations: cMAG — circular MAG, HQ MAG — high-quality MAG, MQ MAG — medium-quality MAG.

To improve MAG recovery from high complexity environmental samples we developed mmlong2, which features multiple optimisations for recovering prokaryotic MAGs from extremely complex metagenomic datasets. Briefly, mmlong2 initially removes eukaryotic contigs, and extracts circular MAGs as separate genome bins (**Figure 1G**). It then performs differential coverage binning (incorporating read mapping information from multi-sample datasets), ensemble binning (using multiple binners on the same metagenome, **Figure S2A**), and iterative binning (metagenome gets binned multiple times iteratively, **Figure S2B**) which all contribute to increased MAG recovery (see **Supplementary Note 1** for more details). In total, 6,076 high-quality (HQ) and 17,767 medium-quality (MQ) MAGs (23,843 total) were recovered by the mmlong2 workflow from the 154 sequenced samples, including 3,349 (14.0 %) MAGs recovered by iterative binning (**Table S2**), with a median of 153.5 (IQR 89.2-203.8) high-or-medium quality MAGs recovered per sample (**Figure 1F**). The obtained MAGs were estimated to account for a median of 24.0 (IQR 16.7-32.9) % of the sequence data within individual samples (**Figure 1E**).

#### 1.2. MAG recovery varies between soil habitats

Lower per-sample sequencing yields were observed from samples originating from the two habitat categories of agricultural fields as well as the bogs, mires and fens habitat (**Figure S3A**), which might be attributed to suboptimal DNA extraction leaving contaminants that compromise the DNA sequencing. In addition, the agricultural field samples had low amounts of sequence data assembled into contigs (median 45.0 %, IQR 39.3-50.1, **Figure S3B**) and also the lowest per-sample count of high-or-medium quality MAGs (median 56.0 MAGs, IQR 34.0-89.0, **Figure S3C**), whereas coastal habitat samples yielded the highest MAG recovery metrics (**Figure S3**). To investigate if the relatively poor MAG yield from agricultural field samples (**Figure S3D**) was only due to sequencing yield, three agricultural and coastal samples were selected and subsampled to specific sequencing depths (from 20 to 100 Gbp, see Methods for more details). Despite normalisation for sequencing effort, the coastal habitat samples still exhibited greater MAG yield (**Figure S4A)**.

Between the two habitat types, there were no significant differences in non-prokaryotic DNA (**Figure S4B**) and overall a comparable number of prokaryotic species was observed in the reads (**Figure S4C**) or contigs (**Figure S4D**), without signs of full microbial diversity capture at 100 Gbp sequencing depth. However, k-mer redundancy analysis indicated that for the coastal samples more species were abundant, compared to the agricultural samples (**Figure S4E-G**). Furthermore, MAGs from the coastal samples also had lower rates of MAG polymorphism (proxy for microdiversity, **Figure S4H**). Hence, the relatively poor MAG recovery from the agricultural field samples was influenced by reduced sequencing yields, higher microdiversity and the absence of dominant species (**Figure S4I**).

### 2. Contribution to the Danish and global environmental microbiomes

#### 2.1. The recovered MAGs are diverse and cover most of Denmark’s core genera

The recovered 23,843 MAGs (**Figure 2A**) were dereplicated into 15,640 different species-level MAGs (**Figure 2B**), comprising 4,894 HQ and 10,746 medium quality (MQ) MAGs (**Figure 2C-F, Figure S5A-B**). Since the MAGs were recovered with long-read Nanopore sequencing, we refer to the dereplicated genome set as the Microflora Danica long-read (MFD-LR) MAG catalogue. The genomic catalogue was inspected for potential Nanopore-associated sequencing errors, and several instances (73, 0.5 % of genomes) of coding density values below 75 % were detected for archaeal MAGs with lower coverage (**Figure S5C**) and reduced guanine-cytosine (GC) content (**Figure S5D**). However, the reduced coding density was also found to be mostly prevalent in archaeal MAGs with increased rates of long homopolymers (more than 6 repeating nucleotides, **Figure S5E**), which in turn were more frequent in MAGs with low GC content (**Figure S5F**).

**Figure 2:**
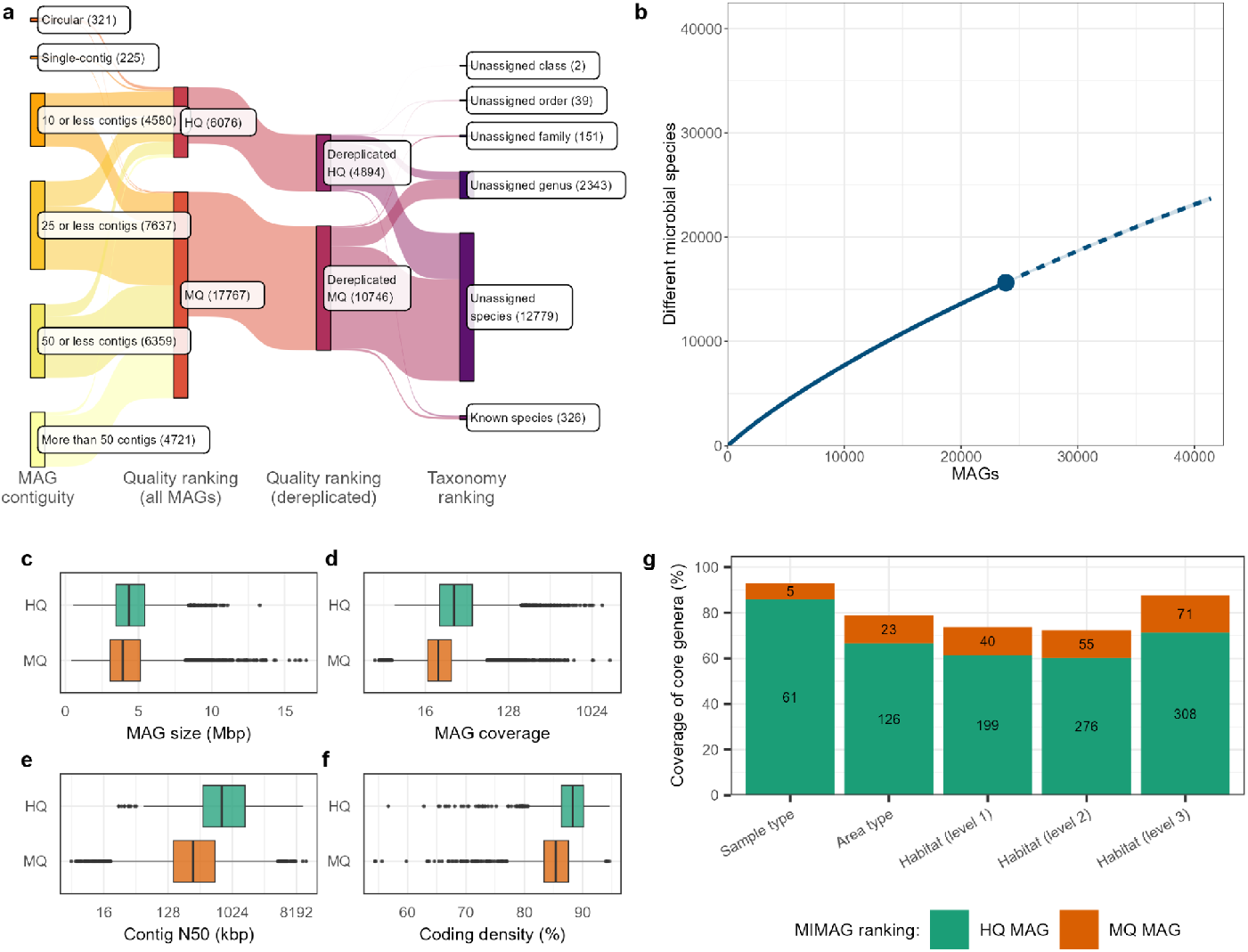
Overview of the MAGs recovered from the sequenced samples. **A)** Aggregated MAG counts by MAG contiguity, MAG quality rankings (MIMAG guidelines) for prior and after MAG dereplication, together with MAG taxonomy rankings, according to GTDB R220 classification. “Single-contig” refers to MAGs that are composed of one contig, which was not reported as circular by the assembler. **B)** Species-level (> 95 % ANI) rarefaction curve for HQ and MQ MAGs recovered in this study. The straight line depicts the rarefaction (interpolation), while the dotted line shows the extrapolation of the curve. **C)** Distribution of dereplicated MAG size values in Mbp, grouped by MAG quality. **D)** Distribution of dereplicated MAG coverage values. **E)** Contig N50 for dereplicated MAGs in kbp. **F)** Coding density values for dereplicated MAGs. **G)** Percentage coverage of core genera of Danish environmental microbiomes^32^ at different sample metadata description levels by the recovered MAGs.

After species-level MAG dereplication, 51.4 % (12,255) of the recovered MAGs were singletons. Plotting a rarefaction curve for the MAGs showcased a near-linear (unsaturated) relationship between the number of recovered MAGs and the number of species-level clusters (**Figure 2B, Figure S6**). The largest species-level cluster of 39 MAGs was recovered for the *Pseudolabrys* genus in the order Rhizobiales, and in total 126 species-level clusters with more than 10 MAGs per cluster were obtained (**Figure S7**).

An advantage of long-read generated MAGs is that they mostly include rRNA operons, enabling direct comparison to the thousands of available 16S rRNA datasets and large-scale databases. Of the recovered dereplicated MAGs, 12,823 (82 %) included at least one 16S rRNA gene and were taxonomically classified against the Microflora Global 16S rRNA database, which features 16S rRNA sequences from the original Microflora Danica project as well as major publicly available 16S rRNA databases^32^. Overall, 12,460 (97.2 %) of these MAGs were classified to the genus level (above 94.5 % 16S rRNA identity), while 10,438 (81.4 %) of the MAGs were assigned a species-level match (above 98.7 % 16S rRNA identity). Coverage of Microflora Danica core genera (across ∼10,000 metagenomes^32^) by the MAG dataset in this study varied from 72.3 % to 93.0 %, depending on habitat descriptor (**Figure 2G**) and was above 90 % for all soil habitats (**Figure S8**).

#### 2.2. Long-read MAGs improve upon previous terrestrial MAG catalogues and boost the characterization of Danish soil microbiomes

Overall, 183 % more dereplicated MAGs were recovered from the 154 deeply sequenced terrestrial samples than the MicroFlora Danica shallow metagenome (MFD-SR) study^45^ that sequenced almost 70 times as many samples – 10,686 samples at ∼5 Gbp each (15,640 vs 5,518 HQ, MQ MAGs), and 10 fold more (4,894 vs 422) dereplicated HQ MAGs were recovered from this study. Also, more MAGs were recovered in this project than the recent genome catalogues of Tibetan Plateau Microbial Catalogue (TPMC^33^) and Old Woman Creek wetland microbial genome catalogue (OWC^34^, **Figure 3A**). Compared to global genomic catalogues that aggregate vast numbers of previously published sequencing data (including low-complexity samples), such as the Searchable, Planetary-scale mIcrobiome REsource (SPIRE^35^), the Genomes from Earth’s Microbiome (GEM^15^), Rare Biosphere Genomes (RBG^36^), and Soil Microbial Dark Matter Metagenome Assembled Genome (SMAG^37^), our 154 long-read sequenced samples still produced similar or higher numbers of HQ genomes, despite, for example, SPIRE utilising almost 100,000 individual samples (**Figure 3A**).

**Figure 3:**
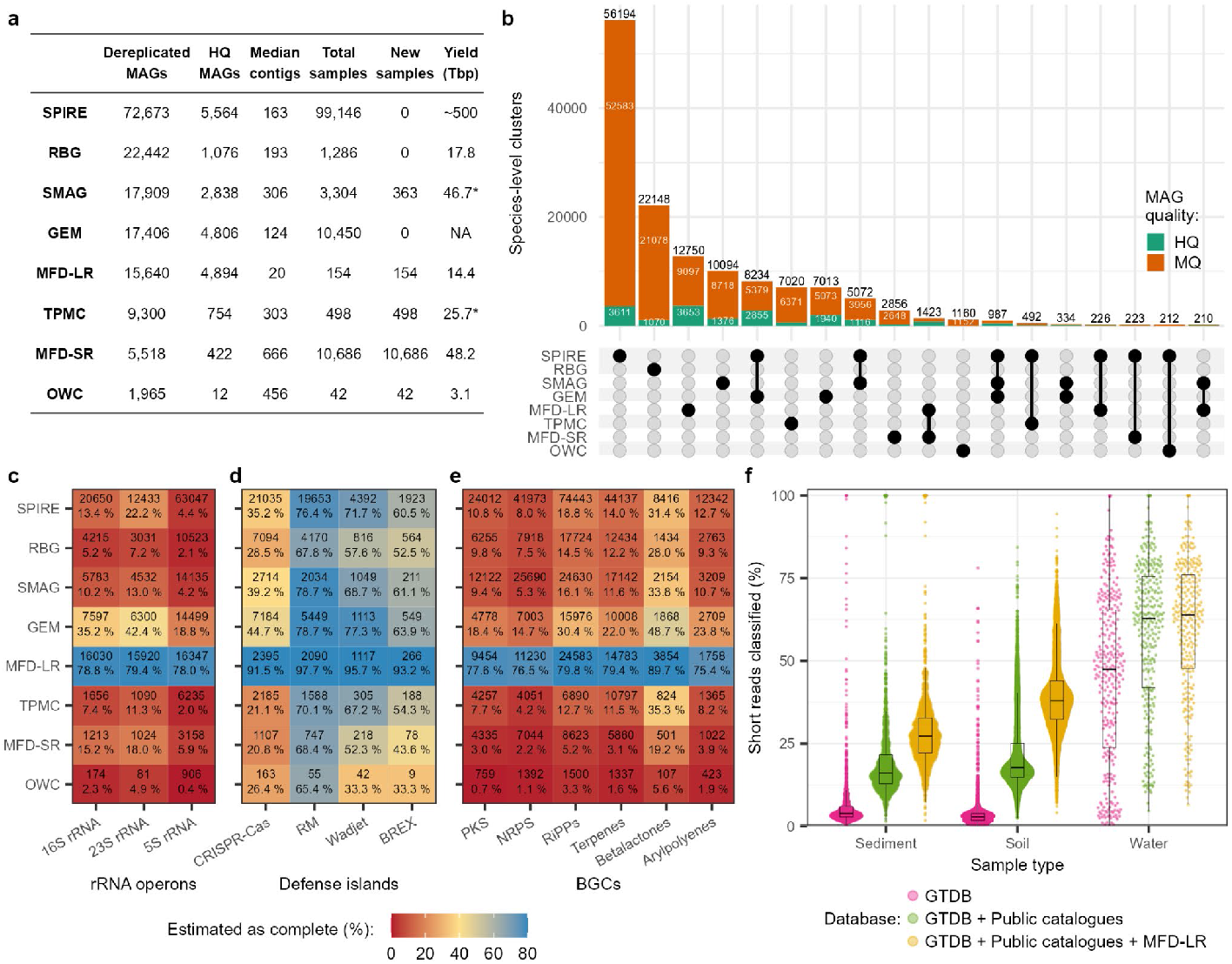
Comparison of MAG catalogues from large-scale terrestrial environment studies. **A)** Catalogue summary, which includes the total number of dereplicated MAGs (HQ and MQ by MIMAG), HQ MAGs, median contig count per dereplicated MAG, number of samples used for MAG recovery, number of samples newly sequenced as part of the study, and the sequencing read data (in Tbp) used. Sequencing yield statistics for the SMAG and TPMC catalogues were extrapolated from the reported read counts, and marked as “NA” for other catalogues when the total sequencing yield was not reported. **B)** Upset plot for species-level MAG overlap between the catalogues. A genome cluster is marked as HQ if at least one of the genomes in the cluster is a HQ MAG. Groups with less than 150 MAGs were omitted from the plot. Total counts of gene clusters for **C)** rRNA operons, **D)** defense islands and **E)** BGCs predicted in the MAGs, grouped by gene cluster type and coloured by the fraction of clusters estimated as complete (see Methods). **F)** Classification rates for Microflora Danica shotgun metagenome datasets using different genome databases, grouped by sample type. “Public catalogues” refers to previously described terrestrial genome catalogues.

Dereplicating MAGs between catalogues resulted in 138,407 species-level clusters, with most species-level overlaps occurring between the reanalysis genome catalogues of SPIRE, SMAG and GEM, due to large overlaps in primary data sources (**Figure 3B**). For MAGs from this study, most species-level overlaps occurred with the short-read MicroFlora Danica MAG catalogue (1,423), although 12,750 dereplicated MAGs (and 3,653 HQ MAGs) from this project represent distinct species.

MAGs from this study also featured greater assembly contiguity with a median contig count of 20 (IQR 10-36) compared to >100 for short-read MAG catalogues (**Figure 3A**). Improved genome contiguity suggests enhanced assembly of complex genomic regions and indeed we observed greatly improved recovery of rRNA genes as part of complete operons (**Figure 3C**) and more complete defense gene islands, especially CRISPR-Cas clusters (**Figure 3D**). More complete biosynthetic gene clusters (BGCs) were also a feature of our improved long read assembly and binning pipeline, and a median of 6.1 fold (IQR: 3.8-14.8) more complete BGCs were observed in the MAGs from this study than other short-read MAG catalogues (**Figure 3E**).

The aforementioned genome catalogues were used as reference databases for classifying the ∼10,000 shallow metagenome datasets from the MicroFlora Danica project^32^. Using the GTDB R220 database alone for read classification resulted in a median species-level classification rate of 3.0 % (IQR 1.9-4.2), whereas including the short-read MAG catalogues increased the median classification rate to 17.3 % (IQR 14.2-25.1). Addition of the long-read MAGs from this study resulted in a database of 229,714 non-redundant genomes and increased species classification to a median of 36.9 % (IQR 30.1-43.3, **Figure 3F**), with the greatest improvements occurring for soil samples (**Figure S9**).

### 3. Novel and expanded microbial lineages

#### 3.1. Long read MAGs expand phylogenetic diversity with novel lineages

Taxonomic classification using GTDB R220 resulted in ANI-based species-level assignments for 326 MAGs, which comprise 2.1 % of the dereplicated MAGs. To determine the novelty and phylogenomic gain for the remaining 15,198 (97.9 %) dereplicated MAGs that could not be assigned a species-level taxonomic label, *de novo* phylogenetic trees were constructed using MAGs from this study and GTDB R220 species representatives (**Figure 4**). MAGs recovered in this study were found to increase the total branch-length of the GTDB prokaryotic genome tree by 8.1 % (**Figure S12**), with most of the branch expansion occurring at genus or species level in both the bacterial and archaeal domains (**Figure S13**). Based on relative evolutionary divergence (RED^38^), this added diversity comprises 1 novel phylum, 21 novel orders, 91 novel families and 1,086 novel genera (**Table S3**).

**Figure 4:**
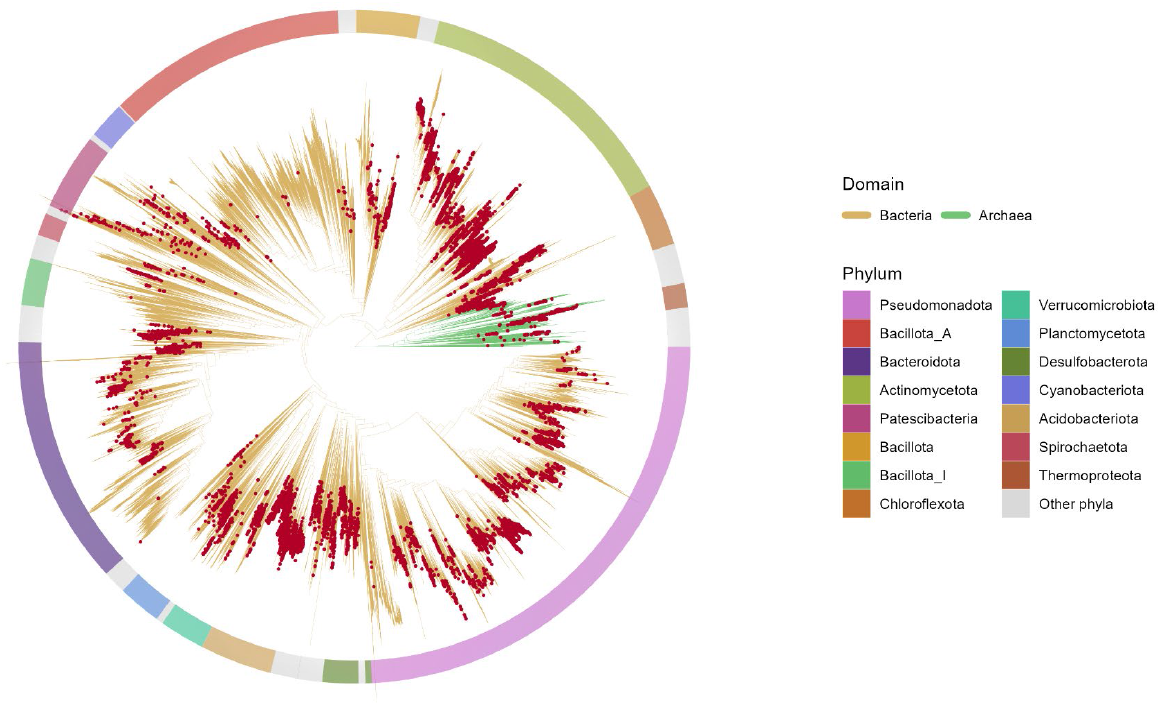
Distribution of the recovered MAGs across the microbial genome tree of life. Microbial genome trees with GTDB R220 representative species for bacteria (120 marker genes) and archaea (53 marker genes) were built separately with 100 bootstraps and merged into a single tree, spanning both domains. Tree tips that represent dereplicated MAGs recovered in this study are marked. The 15 phyla with most genomes are highlighted.

The MAG representing a novel phylum was successfully re-assembled into a circular 2.9 Mbp genome with a GC content of 51.3 %, and a single rRNA operon. The coding density was 91.8 %, although 57.7 % of the predicted genes were hypothetical. Of the genes that could be annotated, the MAG contained multiple genes for key carbon-centric metabolic pathways, such as glycolysis (8/9 complete), pyruvate oxidation, TCA cycle (6/8 complete), ribulose monophosphate pathway, and pentose phosphate pathway (5/6 complete). Across the ∼10,000 Danish environmental metagenomes, species-level matches were observed for 7 samples, 6 of which were from a dystrophic lake habitat, suggesting habitat-specificity and relatively low environmental prevalence of the lineage. Due to the high phylogenetic novelty of the circular MAG, we propose the name *Oederibacterium danicum* sp. nov. in honor of Georg Christian Oeder, a Danish scientist who led the original Flora Danica project^32^.

261 of the novel genera were represented by at least 1 HQ MAG comprising ≤ 10 contigs for which we propose names under the SeqCode^39^. Since the novel MAGs were recovered from Danish habitats, genus names were derived from Danish towns that were nearby the sampling locations and species names derived from environmental features of the samples from which the MAGs were obtained (see Methods). For novel genera that could be assigned to GTDB lineages with placeholder names, we also proposed higher rank names based on the genus stems under the SeqCode to provide taxonomic congruence.

#### 3.2. Long read MAGs greatly expand and provide the first HQ genomes for hundreds of known microbial lineages

MAGs recovered in this study spanned 75 of the 217 currently recognised phyla, with 50 % or higher increases in species-level MAGs for 10 phyla (**Figure S14A, Table S4**). Notably, MAGs were recovered for underrepresented phyla with placeholder names, such as JAUVQV01, CAKKQC01, and UBP4, all of which featured only 2 species in GTDB R220. Furthermore, the highly-populated phyla of Actinomycetota (11,737 representative species in GTDB), Chloroflexota (2,749 species) and Acidobacteriota (1,891 species) have been expanded by 42.1 %, 38.5 % and 134.2 %, respectively, by including novel species-level MAGs obtained in this study. Similar increases in microbial lineage genome counts were observed when examining class, order and family ranks (**Figure S14B-D, Table S4**).

A total of 12,779 dereplicated novel species MAGs were classified as 2,052 different known genera. Of these genera, 682 (32.2 %) were represented by a single MAG in GTDB and inclusion of MAGs recovered in this study expanded the species-level representatives by more than 100 % for 1,065 genera (51.9 % of genera with novel MAGs). The highest number of recovered genomes for a known genus was 320 MAGs for genus Palsa-744 (1,230.8 % increase) belonging to the Actinomycetota, whereas the highest increase in a microbial lineage of 5,000 % was observed for the genus RYN-230 (Actinomycetota) that was represented by a single MAG in GTDB R220.

This study provides the first HQ genomes for 158 known microbial families and 612 known genera (**Table S3**). Notable examples of lineages with newly-added HQ MAGs include the orders of Pacearchaeales (Nanoarchaeota) and Micrarchaeales (Micrarchaeota), which in GTDB R220 are represented by 235 and 189 MQ MAGs, respectively. Similarly, this study provides the first genomes with complete 16S rRNA genes for 436 known genera (**Table S3**), including the Actinomycetota genera *Gaiellasilicea, Gaiella*, and *Desertimonas*, which were all expanded more than 10-fold.

## Discussion

Here we developed mmlong2, a bioinformatic workflow that capitalises on high-throughput deep long-read sequencing to recover MAGs from highly complex terrestrial samples. To evaluate performance, we sequenced 154 soil and sediment samples across 15 environmental habitats. In general, hundreds of MAGs could be recovered from each sample thereby enabling cost-efficient MAG recovery at scale from soils and sediments. However, MAG recovery varied between habitats, especially with agricultural soils consistently yielding fewer MAGs. We show that the variance in MAG recovery between habitats was influenced by sequencing yield, microdiversity and community composition. Hence, we recommend researchers to take into consideration the unique features of each terrestrial habitat when conducting experimental design for future metagenomics projects (e.g. habitat-optimised DNA extraction^40–42^, low-biomass-compatible sequencing protocols^43^). In general, we recommend to sequence at least 60 Gbp per sample, as this ensures access to the genomes of both dominant terrestrial species and low abundance species as evidenced by no indication of saturation observed in the sequencing depth investigated (up to 100 Gbp).

Compared to other large-scale genome-centric studies of terrestrial habitats^15,33,35,37^, this study is the first to use long reads to recover over 10,000 non-redundant MAGs from newly sequenced terrestrial samples. Improved MAG contiguity permits higher resolution of complex genomic regions^44^, such as repeated operons and gene clusters. As the majority of the MAGs were recovered with 16S rRNA genes, most could be linked to the Microflora Global^32^ and other 16S rRNA gene databases. Since rRNA gene databases are generally more diverse than genome databases^45^, recovering more MAGs with complete 16S rRNA genes facilitates improved taxonomic classification and improved linkage between genome and 16S rRNA gene databases^46^. Furthermore, unlike previous terrestrial MAG catalogues, the majority of BGCs and CRISPR-Cas defense islands recovered in this study were estimated to be complete due to improved assembly of the long reads^47–49^ and represents the largest MAG catalogue of complete BGCs to-date, which will enable discovery of medically and industrially valuable biochemical compounds^50^.

Novel HQ and MQ MAGs were recovered for the great majority of genera that were previously reported as constituting the core microbiome of different Danish habitats^32^, thereby enabling further in-depth analysis of functional potential^51^. Also, including MAGs from this study in taxonomically classifying the ∼10,000 short-read Microflora Danica datasets increased median species-level classification from 17.3 to 36.8 %, a significant improvement in the ability to explore complex microbial communities at species level using short-read shotgun metagenomics. The significant improvement in terrestrial metagenome classification also underscores the need for more localised metagenomics projects to acquire genomes of microbes unique to a particular environment or habitat type^52^.

Most of the recovered MAGs from this study constitute novel microbial species or genera, which is a common finding of recent large-scale terrestrial microbiome studies^37,51,53^, highlighting that each genome catalogue contributes significantly to characterising the global microbiome. Although the addition of new microbial lineages from this study occurred mainly at species or genus level, hundreds of recognised order or family level lineages were significantly expanded. As many microbial lineages are currently represented by a single placeholder MAG in GTDB, the expansion of these lineages is imperative to fill the gaps in the tree of life. This study also provides the first HQ MAGs for hundreds of GTDB lineages currently only represented by comparatively fragmented lower quality MAGs. By proposing Latin names for novel lineages under the SeqCode^39^ using HQ MAGs as nomenclatural types we help to address the contemporary issue of an increasingly growing number of unnamed microbial taxa in public databases^54^. As microbial genome databases continuously improve^3^, the quality and not just the quantity of new database additions should be emphasised. Hence, we anticipate this genome catalog will serve as a valuable resource and template for gaining novel insights into the microbial ecology of the world’s most complex environments.

## Methods

### Sample selection

Samples for deep, long-read sequencing were selected using the shallow metagenome OTU tables of 10,686 environmental samples from the Microflora Danica study^32^. Initially, samples with less than 2 Gbp sequencing yield as well as samples from the “Wastewater” habitat were excluded to ensure that the picked samples are from environmental habitats and that the OTU profiles are adequately representative of the sample. Next, OTUs with at least 0.1 % relative abundance and a minimum raw abundance (supporting read count) of 5 were counted and samples, which featured at least 75 of the selected OTUs with a combined relative abundance of 70 %, were further selected to omit samples that are mostly dominated by rare species or belong to a low complexity metagenome. For the remaining samples, OTUs, which were classified as belonging to novel genera within the Microflora Global database^32^ and featured a minimum relative abundance of 0.2 % as well as minimum raw abundance of 10, were counted and 300 samples with the highest number of novel genera OTUs were selected to optimise the likelihood for novel MAG recovery. The remaining samples were then manually curated to optimise for microbial diversity between the samples by omitting samples that overlap based on sampling location or feature high overlapping OTU counts with the rest of the selected samples.

### DNA extraction and Nanopore sequencing

DNA from the selected environmental samples was extracted using the DNeasy PowerSoil Pro Kit (QIAGEN, Germany) and the quality of the extracted DNA was evaluated using the NanoDrop One Spectrophotometer (Thermo Fisher, USA) and the Qubit dsDNA HS kit (Thermo Fisher Scientific, USA, #Q33231) with a Qubit 3.0 fluorometer (Thermo Fisher Scientific, USA) to measure DNA concentration. The DNA was then prepared for sequencing using the SQK-LSK114 Ligation Sequencing kit (Oxford Nanopore Technologies, UK) and loaded into FLO-PRO114M Nanopore flow cells (Oxford Nanopore Technologies, UK) and sequenced in 400 bps sequencing speed mode using either the P2 or the P24 (**Sup dataset 1**) sequencers (Oxford Nanopore Technologies, UK).

### Read data processing

The raw Nanopore sequencing data was collected using the MinKnow software (v22.07.4-23.04.5, **Sup dataset 1**, https://community.nanoporetech.com/downloads) and basecalled with Guppy (v6.2.1-6.5.7, **Sup dataset 1**, https://community.nanoporetech.com/downloads) in super-accurate mode. Due to irreversible updates to the MinKnow software, some samples were sequenced under the 4 kHz sampling rate, while others were acquired using the 5 kHz rate (indicated in **Sup dataset 1**). The sequenced reads were then split with duplex-tools (v0.2.14, https://github.com/nanoporetech/duplex-tools) and trimmed using Porechop (v0.2.3^55^). Reads below Phred Quality score of 7 or length lower than 0.2 kbp were filtered out with NanoFilt (v2.6.0^56^). The split, trimmed and filtered Nanopore read summary statistics were acquired using NanoQ (v0.10.0^57^).

### MAG recovery

MAGs were recovered from the sequenced samples using the custom-developed mmlong2-lite metagenomics workflow v1.0.2 (https://github.com/Serka-M/mmlong2-lite) with the “-sem soil” option used for all samples. The minimum read overlap “-fmc 8” option of the workflow was used with read datasets consisting of more than 50 Gbp of data to speed up the assembly. Multiple shallow metagenome read datasets from the Microflora Danica study^32^ were selected based on overlapping OTU profiles (**Sup dataset 2**) and used as input for the workflow to perform multi-coverage metagenomic binning for improved MAG recovery. Additional information about the mmlong2-lite metagenomics workflow is provided in **Supplementary Note 1**.

### MAG quality control

CheckM1 (v1.2.2^58^) was run on all the recovered MAGs in lineage-specific workflow and all MAGs with less than 50 % completeness or greater than 10 % contamination were omitted. The MAGs were then designated a quality score using CheckM1 metrics with the formula as follows: Genome completeness - Genome contamination * 5. MAGs with a lower quality score than 30 were omitted. The remaining MAGs were then dereplicated using dRep (v2.6.2^59^) with the following settings: “-comp 50”, “-con 10”, “-sa 0.95”, “-nc 0.4”. Furthermore, the MAGs were screened for tRNA molecules with tRNAscan-SE (v2.0.9^60^) using bacterial and archaeal models, while rRNA sequences were detected with Barrnap (v0.9, https://github.com/tseemann/barrnap) and Bakta (v1.9.4^61^), which was run in metagenome mode. Following the minimum information about metagenome-assembled genome guidelines^62^, MAGs were classified into HQ MAGs, if they exhibited greater than 90 % completeness and lower than 5 % contamination estimates by CheckM2 (v1.0.2^63^), while also featuring the 16S, 23S and 5S rRNA genes at least once, together with a minimum of 18 unique tRNAs. MAGs not meeting these criteria, but featuring greater than 50 % completeness and lower than 10 % contamination were classified as MQ MAGs. Only HQ and MQ MAGs, as estimated by both CheckM1 and CheckM2, were used in this study (**Sup dataset 3**).

### Yield-normalised comparisons of environmental habitats

To compare different soil habitats for MAG recovery at normalised sequencing depths, 3 sequenced samples per habitat for agricultural (MFD00392, MFD05176, MFD08497) and coastal (MFD02416, MFD05684, MFD01721) groups were selected and subsampled to custom depths (20, 40, 60, 80, 100 Gbp) with Rasusa (v2.0.0^64^), followed by MAG recovery with mmlong2-lite (v1.0.2). Detection of eukaryotic sequences was performed by classifying the reads and assembled contigs with Kaiju (v1.10.1^65^) using the “kaiju_db_nr_euk_2023-05-10” database. Read and contig taxonomic profiling and species detection was done using Melon (v0.2.0^66^). Variant detection in MAGs for microdiversity assessment was performed with Longshot (v1.0.0^67^). Read K-mer counts were acquired with Jellyfish (v2.2.10^68^), while general read and contig statistics were achieved with Nanoq (v0.10.0^57^) and Cramino (v0.14.1^69^).

### Comparisons to public MAG catalogues

Publically available MAG catalogues were downloaded from the following studies, which featured terrestrial MAGs and at least 1,000 dereplicated MAGs: SPIRE^35^, RBG^36^, GEM^15^, SMAG^37^, TPMC^33^, OWC^34^. MAG quality assessment and quality filtering was performed in the same manner as with the MAGs recovered in this study. For the GEM catalogue, the full MAG dataset was downloaded and dereplication was performed separately to obtain non-redundant MAGs that were recovered in the GEM study. For the SPIRE catalogue, genome entries from the proGenomes database^70^ were omitted and the MAG catalogue was also dereplicated separately, as multiple instances of species-level redundancy were observed for the catalogue. For RBG, MAGs, which were indicated as representative novel species by the authors, were used. The MAGs from all catalogues were then annotated with Bakta (v1.9.4^61^) and screened for defense islands via DefenseFinder (v1.3.0^71^). Screening for secondary metabolites was performed with antiSMASH (v7.1.0^72^) using the following options: “--cb-general”, “--cb-subclusters”, “--cb-knownclusters”, “--genefinding-tool prodigal-m”, “--asf”, “--pfam2go”, “--smcog-trees”, “--rre”, “--tfbs”. The secondary metabolite data was then parsed and aggregated into a dataframe via “tabulate_regions.py” script from multiSMASH (v0.3.0, https://github.com/zreitz/multismash), and gene clusters that were categorized as being coded by the contig edge were considered potentially incomplete. Furthermore, genomes from the catalogues were used for building databases to classify metagenomic reads with Sylph (v0.6.1^73^).

### Phylogenomic analysis

Automated MAG phylogeny and novelty assessment was performed using a custom pipeline available from https://github.com/aaronmussig/mag-phylogeny. Briefly, marker genes were extracted from the MAGs and aligned with the marker genes of GTDB R220 representative genomes using the “infer” module of GTDB-Tk (v2.4.0^74^). The marker gene alignment was then used to build bacterial and archaeal genome trees via FastTree (v2.1.11^75^) with the “WAG” model and 100 bootstraps. RED values for the recovered MAGs was determined with PhyloRank (v0.1.12, https://github.com/dparks1134/PhyloRank) and novelty assignment was performed with “summary_novelty_of_genomes.py” script of the workflow. GTDB representative species classifications, lineage taxonomies and MAG quality rankings were acquired from GTDB R220 metadata files. Phylogenies of highly novel MAGs were manually inspected and metabolic potential was inferred through annotation with DRAM (v.1.4.6^76^).

### Naming of novel microbial lineages

HQ MAGs from novel genera with 10 or less contigs were selected for naming under the SeqCode registry. Latinised genera names were generated by using Danish city or parish names that are within 10 km of the sampling location. Whenever possible, different city or parish names were used to reduce name redundancy. For species names, the species epithet which reflects the sample’s environmental conditions (e.g. sample type or habitat) were given whenever possible. If metadata-based naming was not possible or to reduce name redundancy, generic species names (e.g. danicum, nordicum) were given. Explanations for each of genera and species names are provided in **Sup dataset 4**.

## Data availability

The sequencing read data and the recovered MAGs are available at ENA bio project ID: PRJEB58634. Code, datasets used for plotting the figures and other resources are available at https://github.com/Serka-M/mfd_mags. MAG recovery workflow used in the study is available at https://github.com/Serka-M/mmlong2-lite. Yield-normalised comparative metagenomics workflow is available at https://github.com/Serka-M/mmcomp. MAG phylogeny workflow is available at https://github.com/aaronmussig/mag-phylogeny. GTDB-tk database used for MAG taxonomy can be accessed at https://data.ace.uq.edu.au/public/gtdb/data/releases/release220. The Microflora Global 16S rRNA database is available at https://doi.org/10.6084/m9.figshare.26030524.v2. Different MAG catalogues used in this study is available as follows: MFD-SR — https://zenodo.org/records/12605769, SPIRE — https://spire.embl.de/downloads, TPMC — https://download.cncb.ac.cn/bigd/TPMC/, RBG — https://connectqutedu.sharepoint.com/:f:/s/BinChickensupplementarydata/ErJbGzlvzglMoLSAIjv3dtIBaEmqdZXoUHYRQlYYbtSY5Q?e=aIvFdH, OWC — https://zenodo.org/records/8194033, SMAG — https://bma-public.s3.cn-northwest-1.amazonaws.com.cn/SMAG/magdrep.tar.gz, GEM — https://portal.nersc.gov/GEM.

## Acknowledgments

This study was funded by a research grant from Poul Due Jensen Foundation (Microflora Danica) and VILLUM FONDEN (130690, 50093). We would like to thank the Microflora Danica Consortium for their contributions to sample and metadata collection across Denmark.

## Author Contributions Statement

MA, PHN, MS designed the study. TBNJ, EAS, YY contributed to the generation or processing of the short-read shotgun metagenome or 16S amplicon sequencing data used in the study. VRJ, FD performed curation and validation of the sample metadata. MS, CMS, AKSK, FP performed sample selection for sequencing. MS and CJ performed sample DNA extraction and Nanopore sequencing. MS carried out the long-read sequencing data processing, MAG recovery, MAG analysis, and writing of the initial manuscript. AJM, PH, MS performed phylogenetic analysis of the MAGs. All authors reviewed the manuscript.

