## Supplementary material for "Recovery of highly contiguous genomes from complex terrestrial habitats reveals over 15,000 novel prokaryotic species and expands characterization of soil and sediment microbial communities"

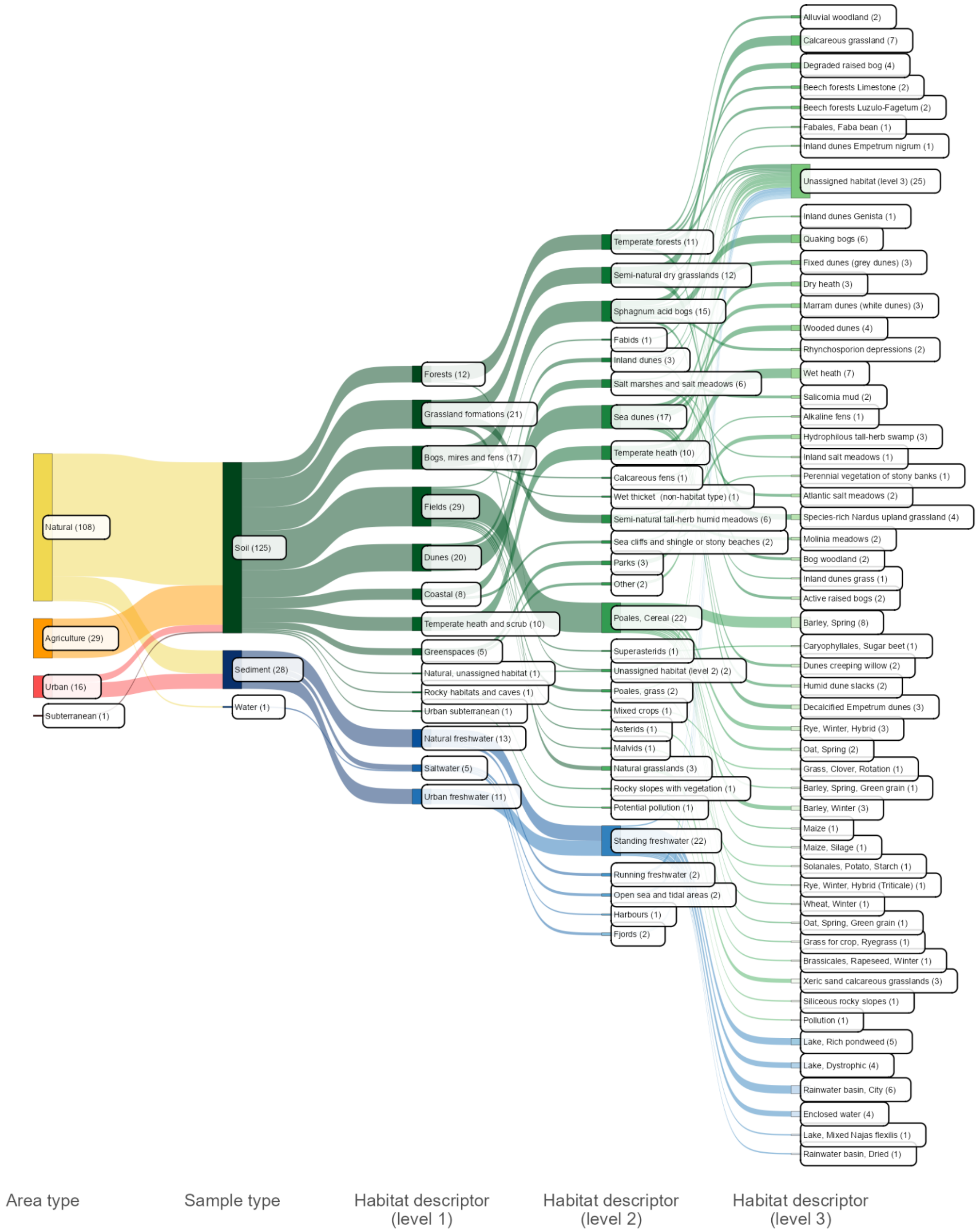


**Figure S1:** Overview of the sequenced sample habitat ontology. The presented sample metadata is derived from the Microflora Danica 10 000 shallow metagenome database.

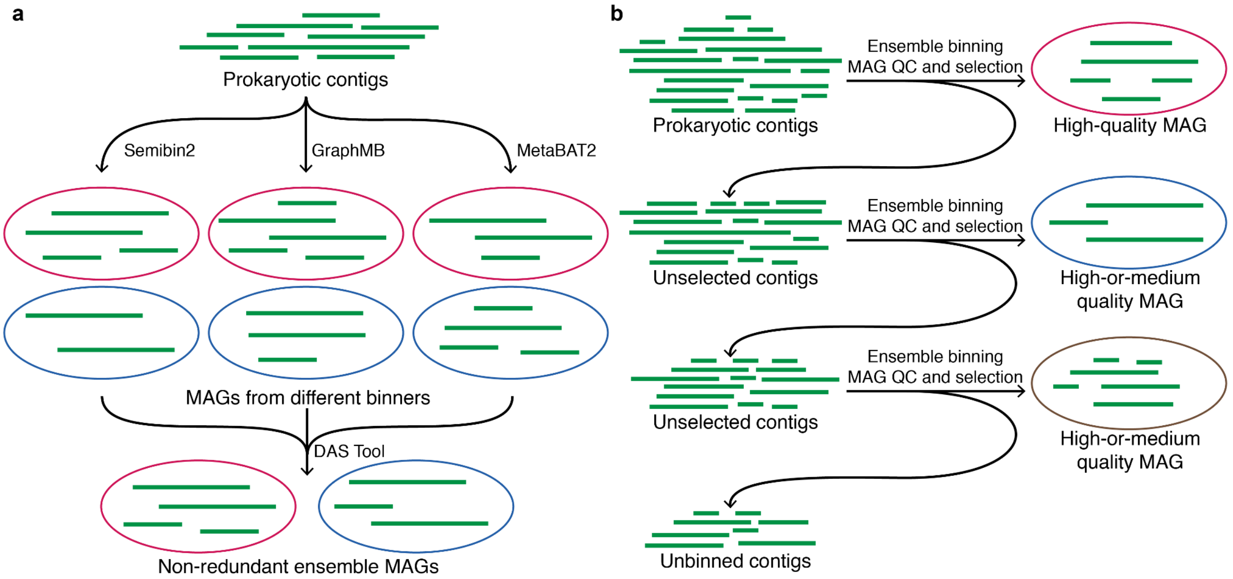


**Figure S2:** Schematic overview of the key binning procedures used by mmlong2 metagenomics workflow. **A)** Illustrative overview of the ensemble binning process used by mmlong2 for MAG production. **B)** Illustrative overview of the iterative metagenomic binning process by mmlong2.

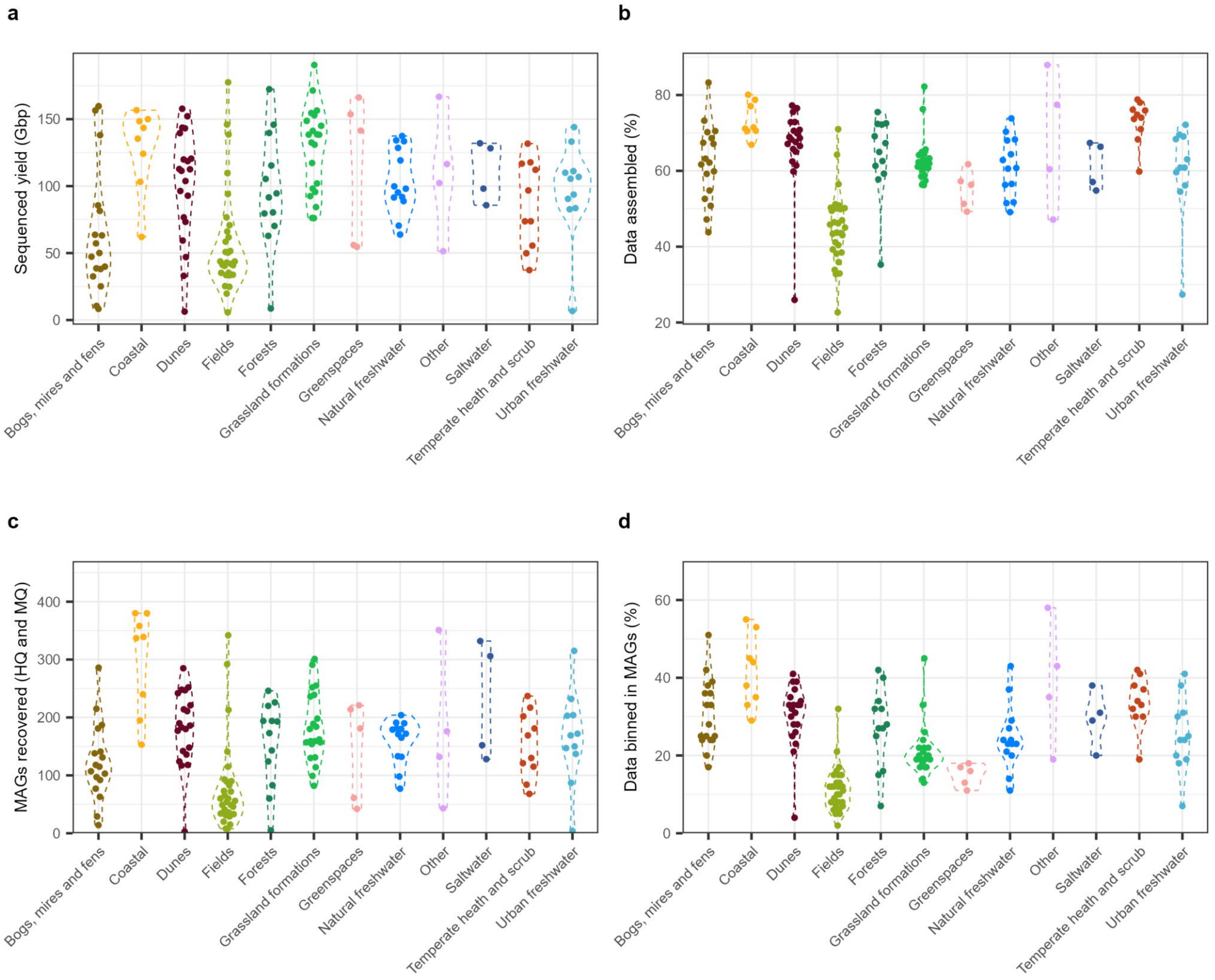


**Figure S3:** Per-sample sequencing and metagenome assembly metrics, grouped and colored by sample habitat. **A)** Sequenced data yield in Gbp. **B)** Percentage values for sequenced read data that was assembled into metagenomic contigs. **C)** HQ and MQ MAGs recovered. **D)** Per-sample percentage of sequenced data accounted for by HQ and MQ MAGs. Habitats represented by a single sample were grouped into the "Other" category.

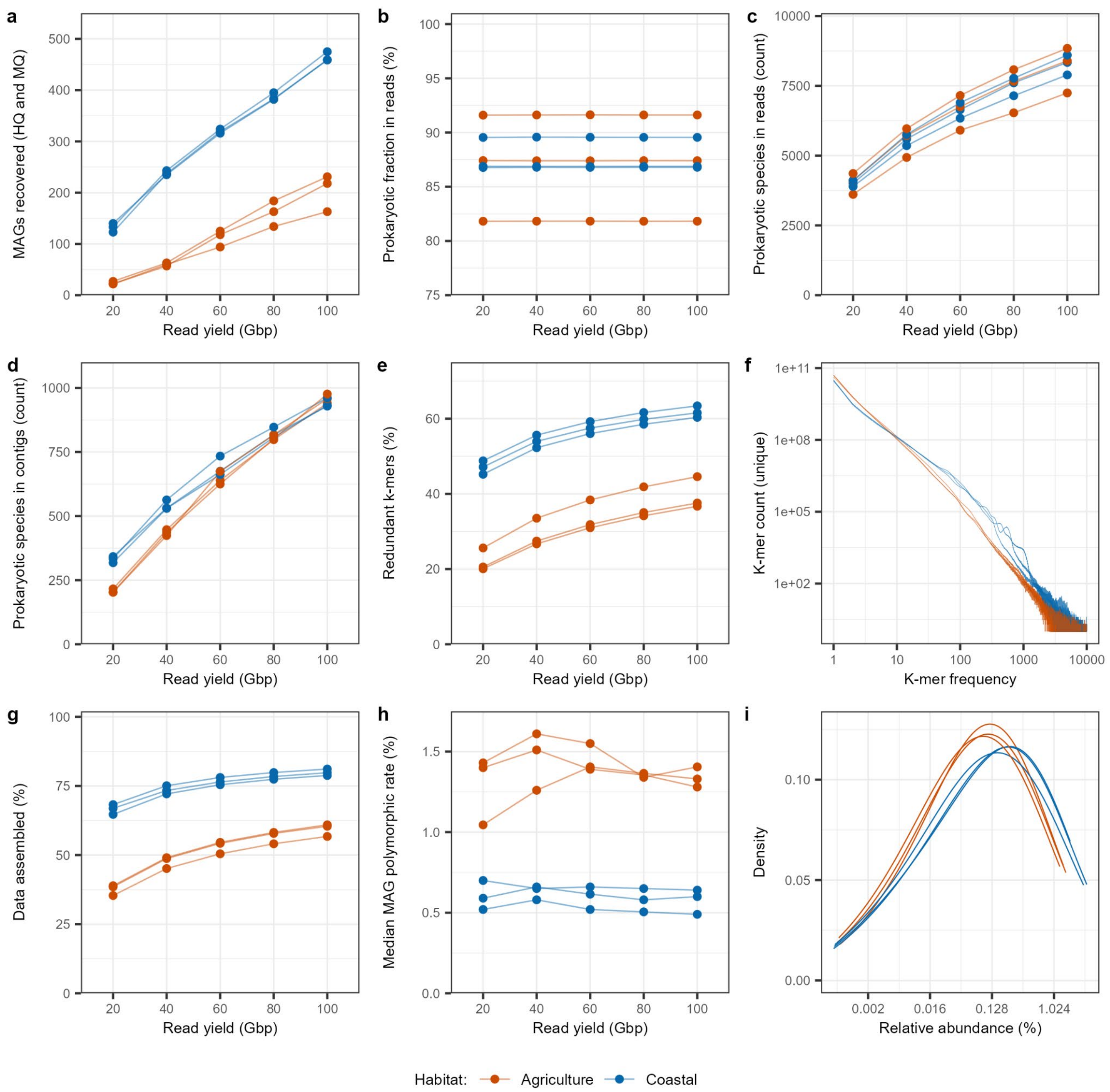


**Figure S4:** Comparison of coastal and agricultural habitats for MAG recovery at normalised sequencing depths. 3 samples were used per habitat. **A)** Recovered HQ and MQ MAG counts. **B)** Rates of sequenced reads classified as prokaryotic. **C)** Counts for observed prokaryotic species with Melon taxonomic profiler in read datasets. **D)** Number of prokaryotic species observed in the assembled metagenomes with Melon taxonomic profiler. **E)** Rates for k-mer (24-mer) redundancy. **F)** K-mer (24-mer) spectrum plots for 100 Gbp datasets. **G)** Rates of sequenced reads assembled into contigs. **H)** Median recovered MAG polymorphism rates. **I)** Density curves for prokaryotic species relative abundances (log-scaled) within the 100 Gbp read datasets, as observed with Melon taxonomic profiler.

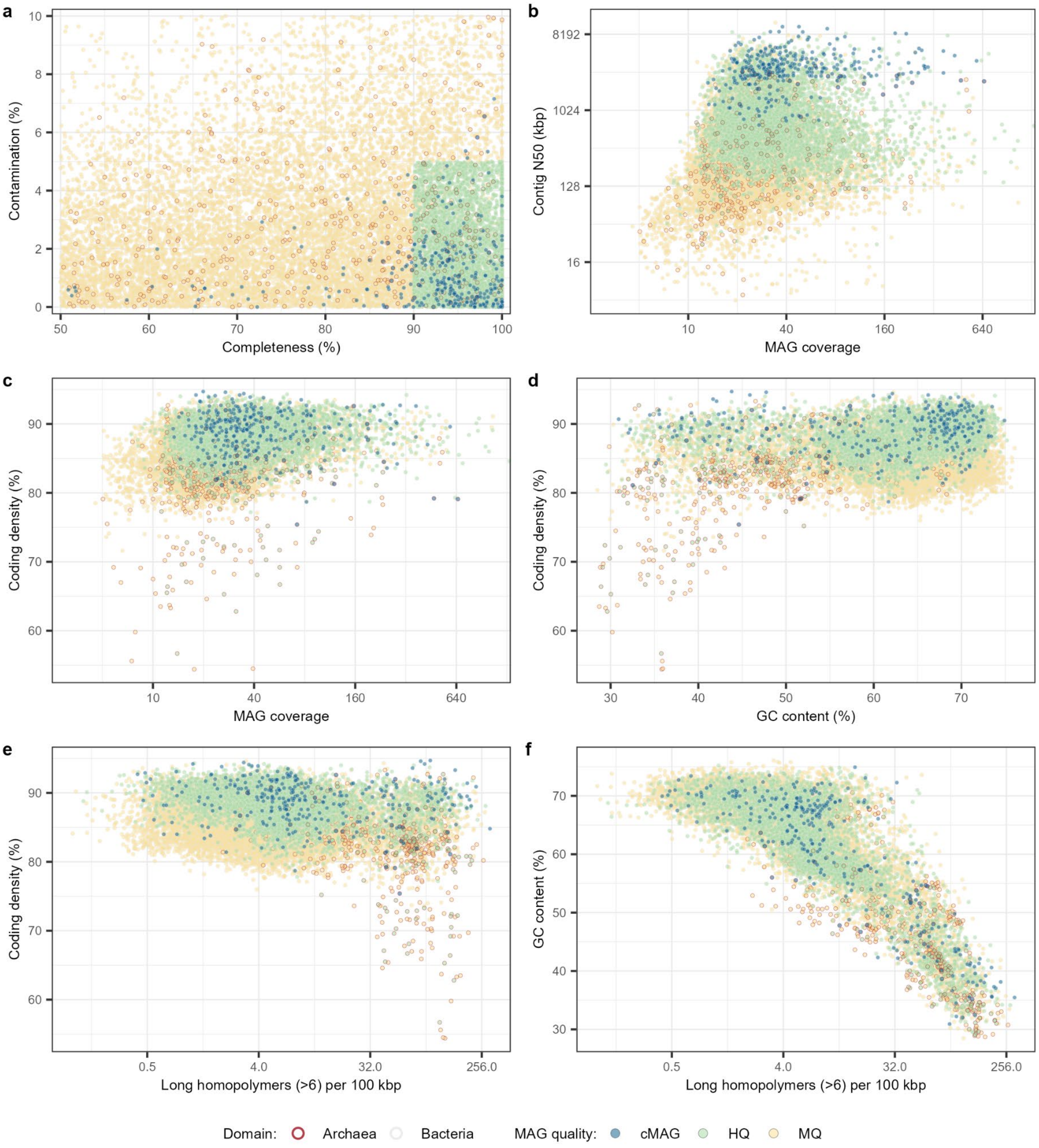


**Figure S5:** General quality metrics for dereplicated MAGs recovered in this study, coloured by MAG quality. **A)** MAG completeness and contamination values by CheckM2. **B)** MAG contig N50 values by MAG coverage. **C)** Coding density values by MAG coverage. **D)** Coding density values by MAG GC content. **E)** Coding density values by long homopolymer counts. **F)** MAG GC content by long homopolymer counts.

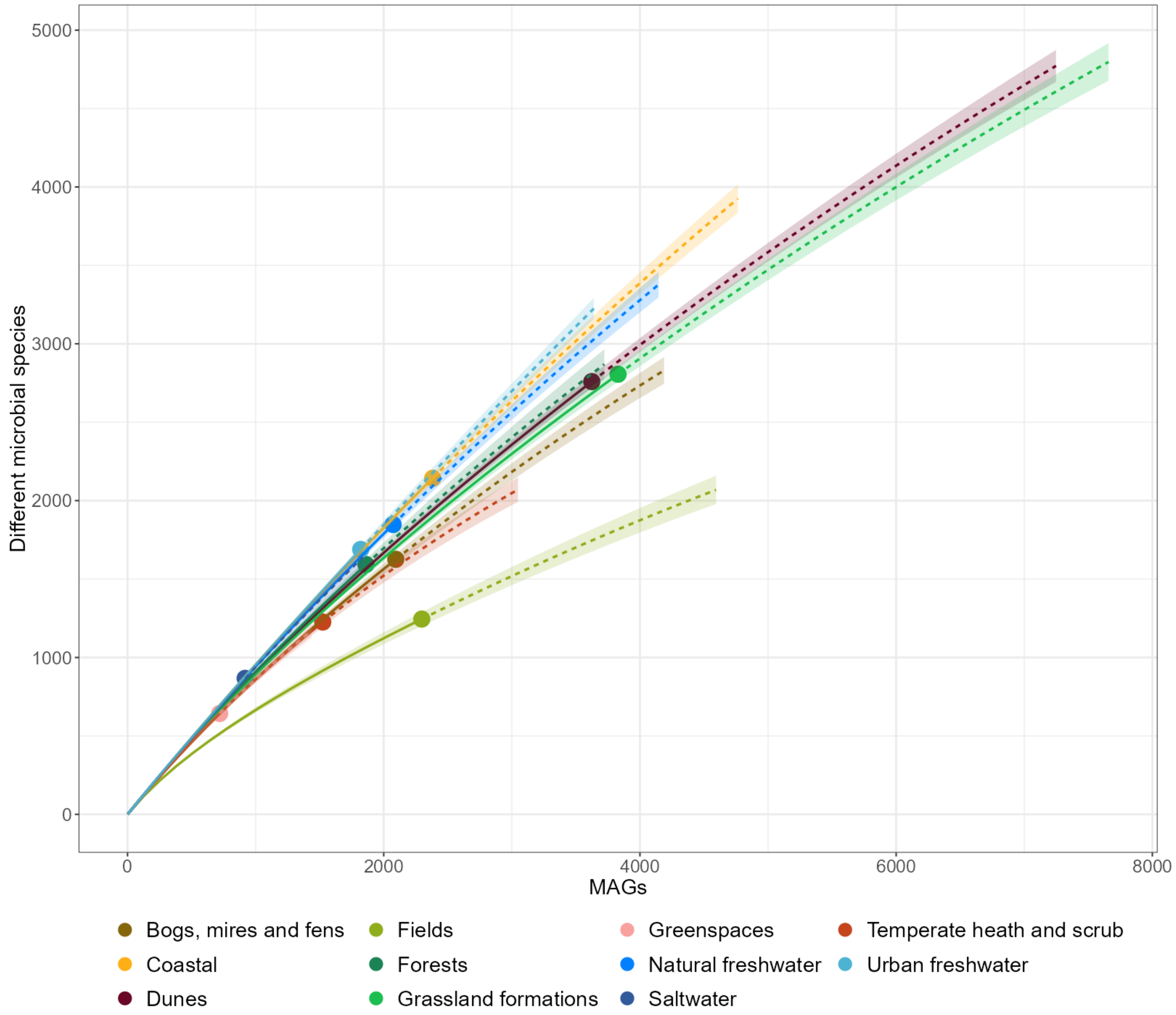


**Figure S6:** Species-level MAG rarefaction curves for HQ and MQ MAGs recovered in this study, grouped and colored by sample habitat. The straight line shows the rarefaction (interpolation), while the dotted line indicates the extrapolation of the curve. Habitats represented by a single sequenced sample were omitted from the plot.

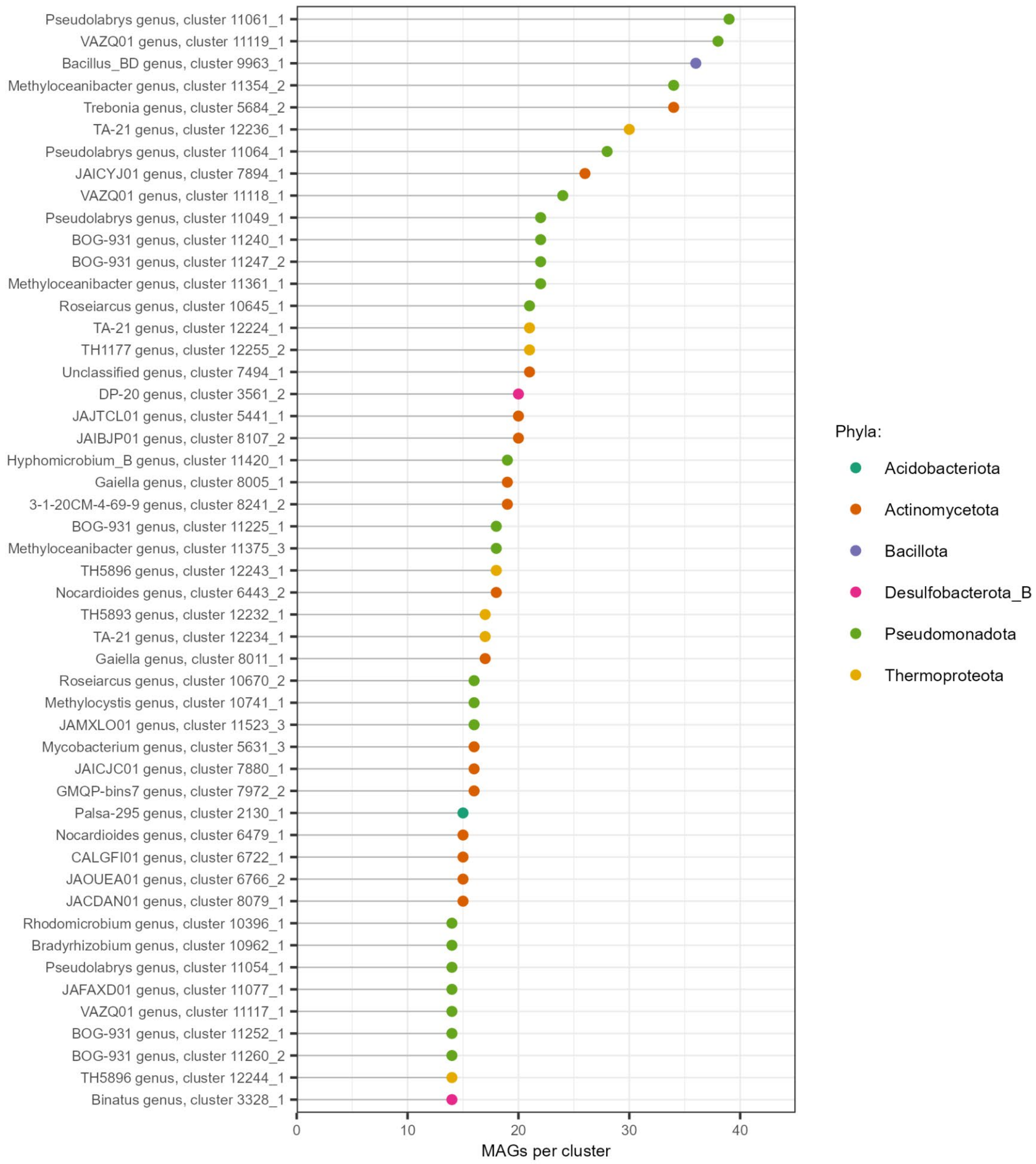


**Figure S7:** Top 50 largest species-level clusters acquired after MAG dereplication, coloured by phylum. Genus (if classified) and corresponding species-level cluster ID is provided.

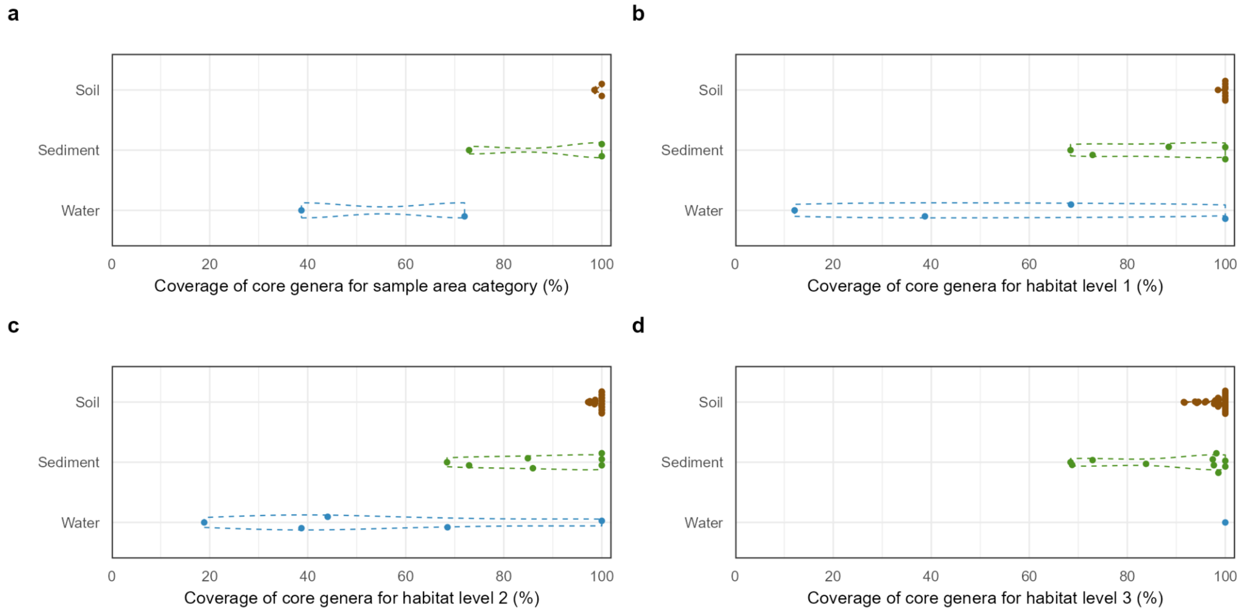


**Figure S8:** Coverage of the core genera of Danish environmental microbiomes by the recovered long-read MAGs. The coverage values are grouped by sample type and presented for the following Microflora Danica metadata categories: **A)** area type, **B)** habitat descriptor level 1, **C)** habitat descriptor level 2, **D)** habitat descriptor level 3. Only the dereplicated MAGs with 16S rRNA sequences that could be classified above 94.5 % sequence identity were included in the calculations.

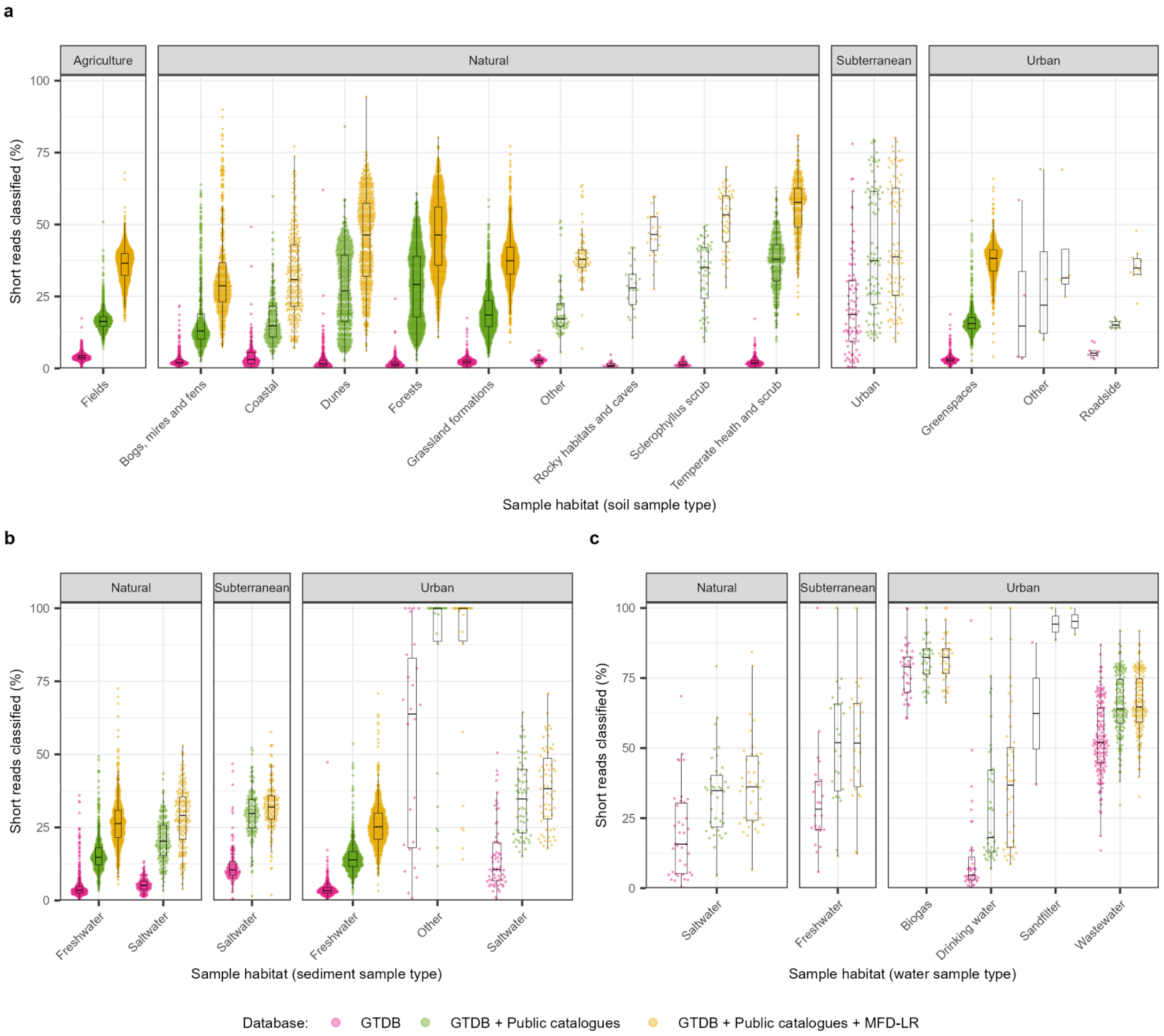


**Figure S9:** Classification rates for Microflora Danica shallow metagenome datasets using different genome databases, grouped by sample habitat. Read classification rates are presented for sample types of **A)** soil, **B)** sediment, and **C)** water samples. The samples are further faceted by sample area type. “Public catalogues” refers to previously described terrestrial genome catalogues.

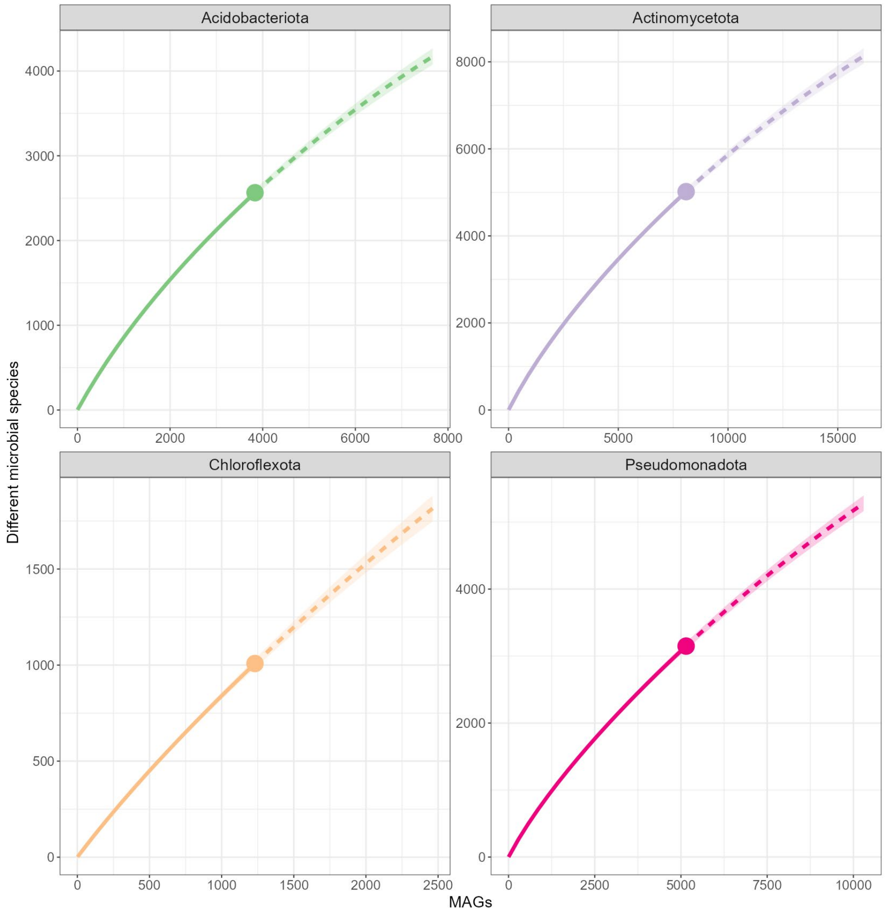


**Figure S10:** Species-level MAG rarefaction curves for HQ and MQ MAGs recovered in this study, provided for the most frequent 4 phyla. The straight line shows the rarefaction (interpolation), while the dotted line indicates the extrapolation of the curve.

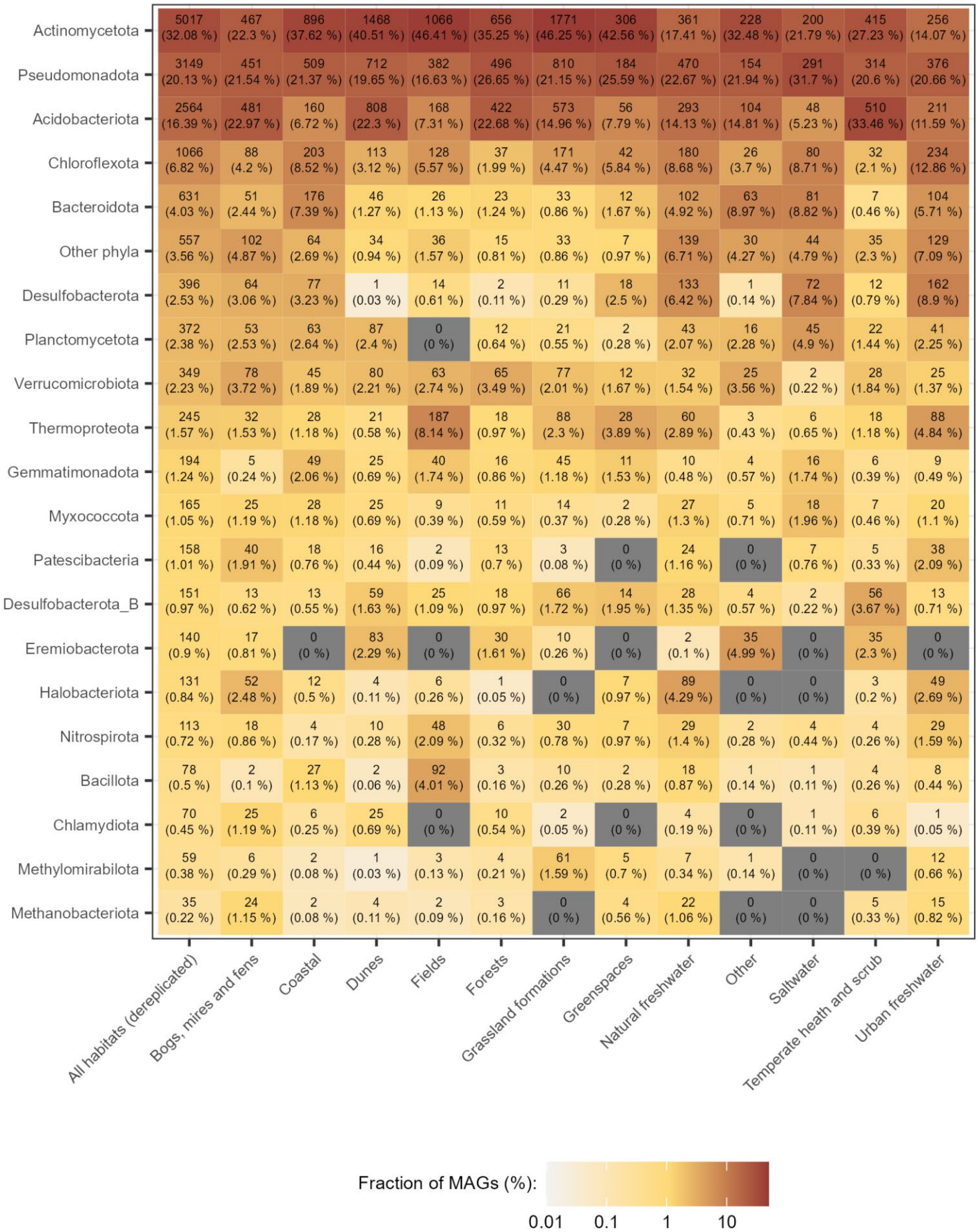


**Figure S11:** MAG counts and percent fractions by phylum-level MAG taxonomic classification via GTDB-tk, grouped by habitat. The "All habitats" column refers to aggregated and dereplicated MAG taxonomy counts from all samples in this study.

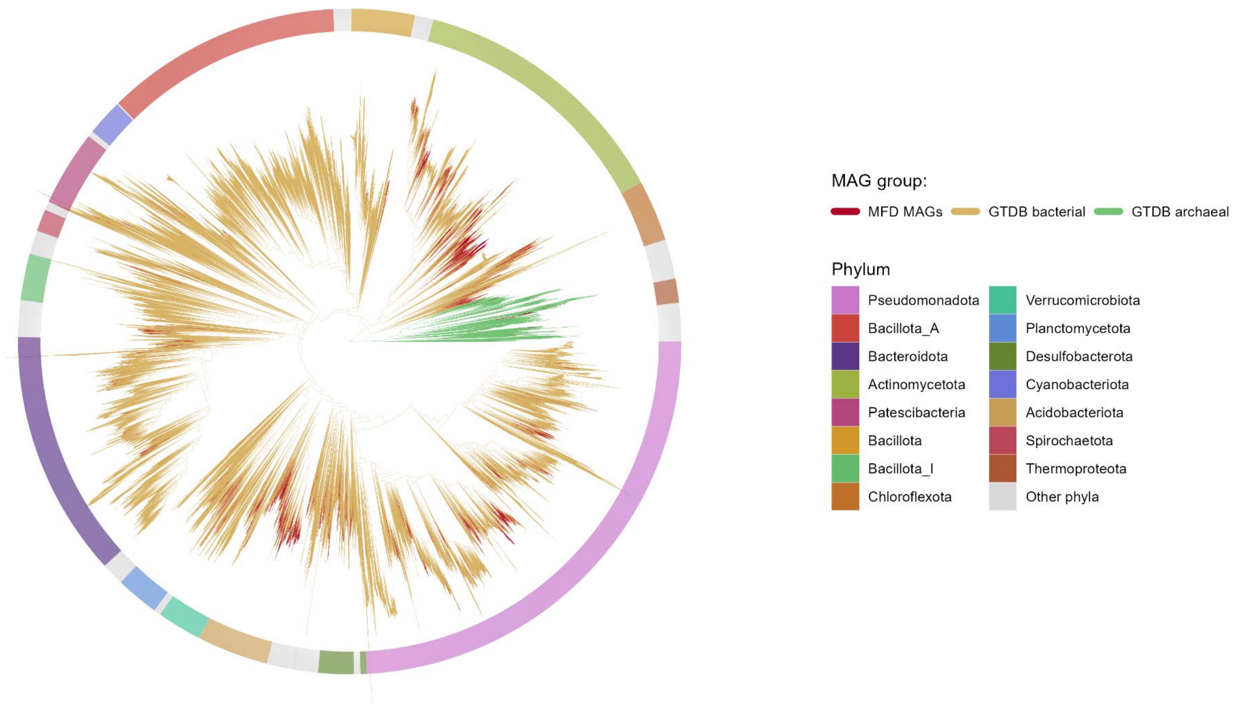


**Figure S12:** Contribution of MAGs recovered in this study to the microbial tree of life. Microbial genome trees with GTDB R220 representative species for bacteria (120 marker genes) and archaea (53 marker genes) were built separately with 100 bootstraps and merged into a single tree, spanning both domains. Branches, which are constituted by MAGs recovered in this study are highlighted. The top 15 most common phyla are highlighted.

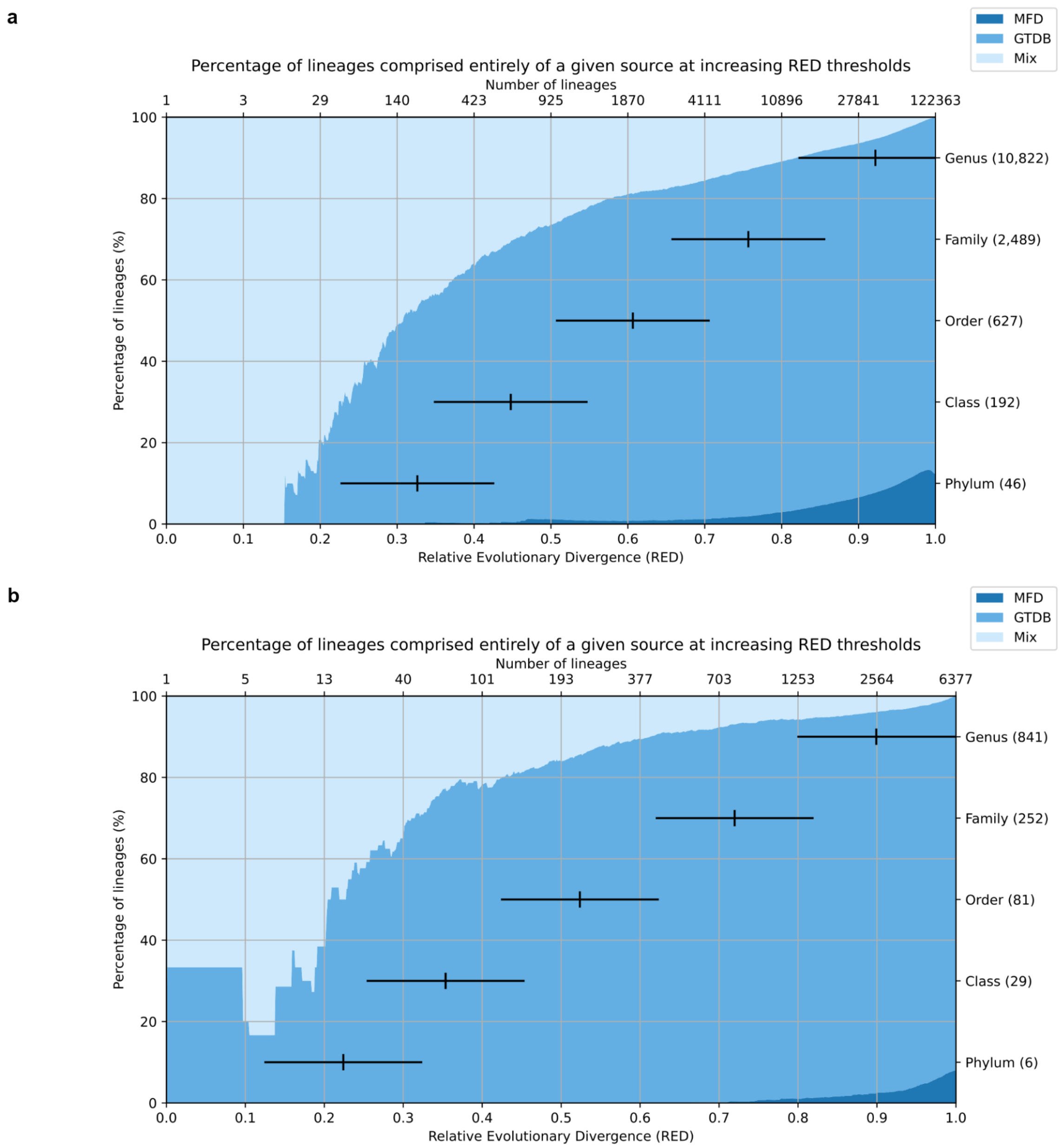


**Figure S13:** Summary of phylogenetic gain to GTDB R220 from novel lineages at different phylogenetic ranks. The summaries are presented for bacterial (**A**) and archaeal (**B**) domains.

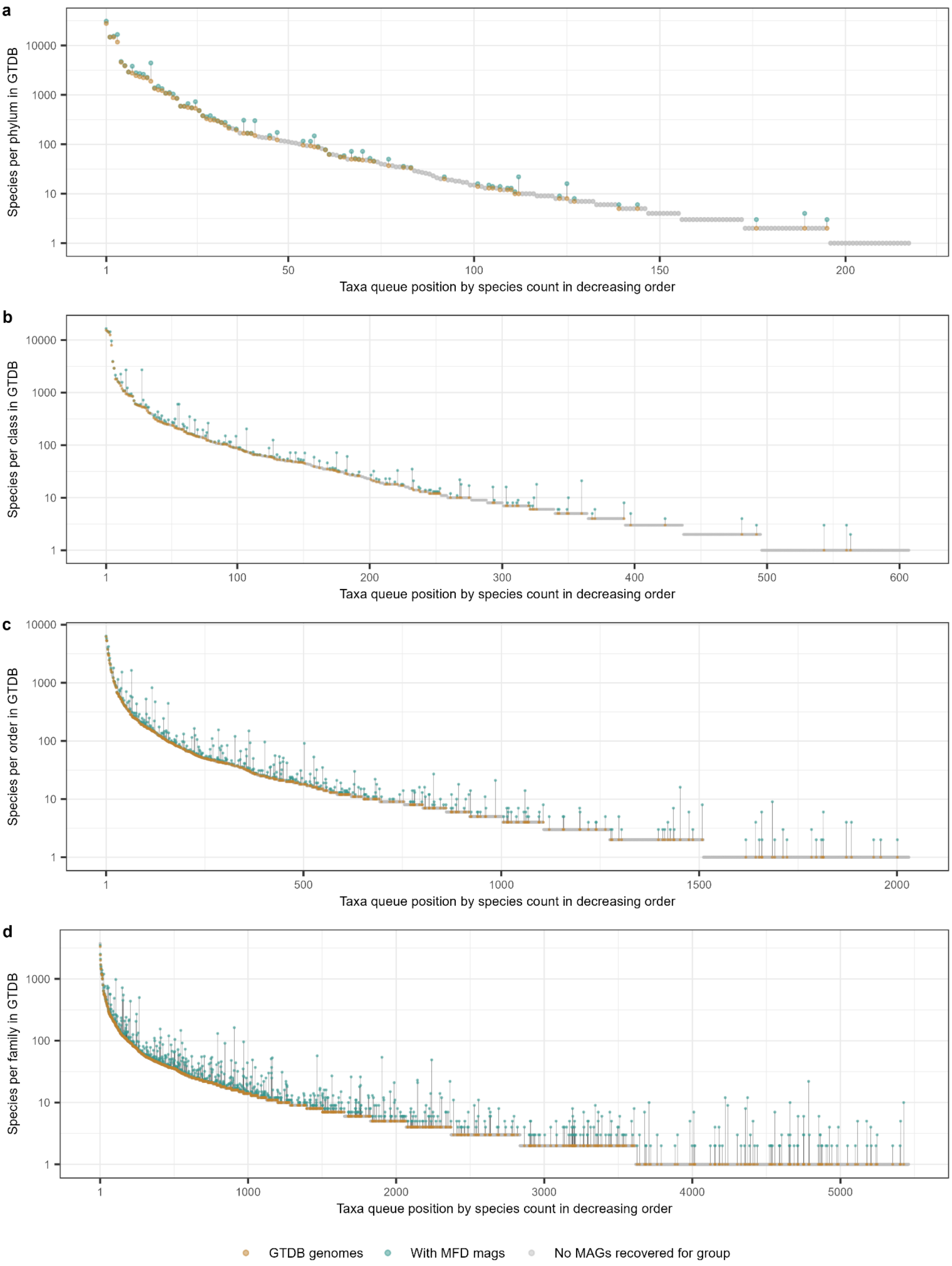


**Figure S14:** Increase in species-level genomes for GTDB R220 microbial lineages after including novel MAGs recovered in this study. The increases in genome counts per microbial lineage are provided for **A)** phylum, **B)** class, **C)** order, and **D)** family ranks.

**Table S1:** Habitat-level overview of the sequenced samples.

| Sample type | Sample area | Sample habitat | Samples | Yield, Gbp | Total MAGs | HQ MAGs |
|---|---|---|---|---|---|---|
| Soil | Natural | Grassland formations | 21 | 2,681.9 | 3,829 | 769 |
| Soil | Natural | Dunes | 20 | 2,020.7 | 3,624 | 1,109 |
| Soil | Natural | Bogs, mires and fens | 17 | 1,095.8 | 2,094 | 496 |
| Soil | Natural | Forests | 12 | 1,166.4 | 1,861 | 475 |
| Soil | Natural | Temperate heath and scrub | 10 | 865.4 | 1,524 | 479 |
| Soil | Natural | Coastal | 8 | 1,023.5 | 2,382 | 850 |
| Soil | Natural | Rocky habitats and caves | 1 | 166.7 | 351 | 112 |
| Soil | Natural | Unassigned habitat | 1 | 116.4 | 132 | 27 |
| Soil | Agriculture | Fields | 29 | 1,694.3 | 2,297 | 293 |
| Soil | Urban | Greenspaces | 5 | 571.9 | 719 | 81 |
| Soil | Subterranean | Urban subterranean | 1 | 51.4 | 43 | 12 |
| Sediment | Natural | Saltwater | 4 | 444.1 | 918 | 315 |
| Sediment | Natural | Freshwater | 13 | 1352.5 | 2,073 | 468 |
| Sediment | Urban | Freshwater | 11 | 1,068.0 | 1,820 | 492 |
| Water | Natural | Saltwater | 1 | 102.3 | 176 | 98 |
| | | **Total** | 154 | 14.42 | 23,843 | 6,076 |

**Table S2:** Summary of MAG recovery by the mmlong2 workflow.

| Binning stage | Total MAGs | HQ MAGs | Total singleton MAGs | Singleton HQ MAGs |
|---|---|---|---|---|
| cMAG extraction | 321<br>(1.3 %) | 247<br>(4.1 %) | 258<br>(2.1 %) | 194<br>(6.1 %) |
| Round 1 | 6,088<br>(25.5 %) | 5,340<br>(87.9 %) | 3,075<br>(25.1 %) | 2,772<br>(87.1 %) |
| Round 2 | 14,449<br>(60.6 %) | 364<br>(6.0 %) | 7,311<br>(59.7 %) | 158<br>(5.0 %) |
| Round 3 | 2,603<br>(10.9 %) | 120<br>(2.0 %) | 1,409<br>(11.5 %) | 58<br>(1.8 %) |
| Round 4 | 382<br>(1.6 %) | 5<br>(0.1 %) | 202<br>(1.6 %) | 2<br>(0.1 %) |
| **Total** | 23,843 | 6,076 | 12,255 | 3,184 |

**Table S3:** Summary for the contribution of the recovered MAGs to GTDB R220.

| Rank | Lineages in GTDB R220 | Novel lineages recovered | Lineages without HQ MAGs* | First HQ MAGs recovered | Lineages without 16S rRNA* | First MAGs with 16S rRNA recovered |
|---|---|---|---|---|---|---|
| Phylum | 217 | 1 | 72 | 2 | 26 | 1 |
| Class | 607 | 1 | 277 | 8 | 86 | 3 |
| Order | 2,030 | 21 | 1,142 | 46 | 324 | 12 |
| Family | 5,460 | 91 | 3,581 | 158 | 1,095 | 54 |
| Genus | 24,959 | 1,086 | 18,228 | 612 | 7,398 | 436 |
| Species | 113,104 | 15,314 | 85,812 | 4,757 | 61,767 | 12,541 |

*Refers to the lineages in GTDB R220

**Table S4:** Summary of GTDB R220 prokaryotic lineage expansion by the recovered MAGs.

| **Rank** | **Lineages in GTDB** | **Expansion by ≥ 1 MAG** | **Expansion of ≥ 25 %** | **Expansion of ≥ 50 %** | **Expansion of ≥ 100 %** |
|---|---|---|---|---|---|
| Phylum | 217 | 75 | 16 | 10 | 4 |
| Class | 607 | 182 | 61 | 38 | 22 |
| Order | 2,030 | 451 | 183 | 134 | 69 |
| Family | 5,460 | 864 | 480 | 354 | 225 |
| Genus | 24,959 | 2,052 | 1,649 | 1,358 | 1,065 |

**Supplementary Note 1:** Brief description of mmlong2-lite metagenomics workflow.

The mmlong2-lite metagenomics workflow v1.0.2 is a Snakemake[1] bioinformatics workflow that can take long reads (Nanopore or PacBio HiFi) and perform metagenome assembly, contig filtering, binning and initial MAG quality check. For Nanopore datasets, the reads are assembled into metagenomes using Flye (v2.9.2[2]) with "--meta" and "--nano-hq" options. Furthermore, the "-fmc" flag of mmlong2-lite controls the "min_read_cov_cutoff" option of Flye, which can be increased to filter out more low coverage contigs and thus speed up the metagenome assembly turnaround time.

The assembled Nanopore-only metagenome is then polished with 1 round of Medaka (v1.8.0, https://github.com/nanoporetech/medaka) to reduce the amount of indel errors in the initial assembly. Contigs shorter than 3 kbp are filtered out using SeqKit (v2.4.0[3]) and the remaining contigs are then classified with Tiara (v1.0.3[4]) to remove eukaryotic contigs from the assembly.

Prior to metagenomic binning, the "assembly_info.txt" file outputted by Flye is used to extract circular contigs above 250 kbp from the assembly and placed into separate bins. The remaining linear contigs are used for iterative ensemble binning with MetaBAT2 (v2.15[5]), SemiBin2 (v1.5[6]), GraphMB (v0.1.5[7]), and DAS Tool (v1.1.3[8] and the "--search_engine diamond" setting). If provided with the "-cov" option, contig multi-coverage profiles are generated from different read datasets (long or short reads) by mapping the sequenced reads to the metagenome using Minimap2 (v2.26[9]) and SAMtools (v1.16.1[10]), followed by coverage calculation using the "jgi_summarize_bam_contig_depths" function of MetaBAT2. The concatenated coverage profiles are then provided as input to MetaBAT2 and GraphMB, while for SemiBin2, the mapping files are provided directly.

After acquiring the ensemble bins with DAS Tool, CheckM2 (v1.0.2[11]) is used to acquire bin completeness and contamination metrics, followed by selection of bins meeting the requirements for HQ MAGs (> 90 % completeness, < 5 % contamination). The unselected contigs are then binned again using the same binners and then all HQ as well as MQ MAGs (> 50 % completeness, < 10 % contamination) are selected. Ensemble binning with the unselected contigs is repeated for the third time with the ensemble MAG filtering feature of DAS Tool turned off as well as the use of a pre-trained binning model for SemiBin2 ("global" model by default), followed by retainment of all HQ and MQ MAGs. The remaining unselected contigs are then binned one last time with MetaBAT2 and only HQ as well as MQ MAGs, as estimated by CheckM2, are kept as the final output of the MAG production workflow.

MAG relative abundance and coverage values are computed using CoverM (v0.6.1, https://github.com/wwood/CoverM), while Quast (v5.2.0[12]) is also run on the MAGs to acquire quality metrics. The mmlong2-lite workflow is publicly available at https://github.com/Serka-M/mmlong2-lite as well as https://zenodo.org/record/8013498.
